# Structures of the holo CRISPR RNA-guided transposon integration complex

**DOI:** 10.1101/2022.10.12.511933

**Authors:** Jung-Un Park, Amy Wei-Lun Tsai, Alexandrea N. Rizo, Vinh H. Truong, Tristan X. Wellner, Richard D. Schargel, Elizabeth H. Kellogg

**Affiliations:** Department of Molecular Biology and Genetics, Cornell University, Ithaca, NY 14853, USA.

## Abstract

CRISPR-associated transposons (CAST) are programmable mobile elements that insert large DNA cargo by an RNA-guided mechanism. Multiple conserved components act in concert at the target site through formation of an integration complex (transpososome). We reconstituted the type V-K CAST transpososome from *Scytonema hofmannii* (ShCAST) and determined the structure using cryo-EM. Transpososome architecture ensures orientation-specific association: AAA+ regulator TnsC has defined polarity and length, with dedicated interaction interfaces for other CAST components. Interestingly, transposase (TnsB)-TnsC interactions we observe contribute to stimulating TnsC’s ATP hydrolysis activity. TnsC deviates from previously observed helical configurations of TnsC, and target DNA does not track with TnsC protomers. Consequently, TnsC makes new, functionally important protein-DNA interactions throughout the transpososome. Finally, two distinct transpososome populations suggests that associations with the CRISPR effector are flexible. These ShCAST transpososome structures significantly enhances our understanding of CAST transposition systems and suggests avenues for improving CAST transposition for precision genome-editing applications.

## Introduction

CRISPR-associated transposons (CAST) are Tn7-like mobile elements (*1*) that programmably integrate large DNA cargo at genomic locations dictated by guide-RNA sequence (*2, 3*). Multiple independent domestication events (*1, 4*) created the distinct CAST sub-families characterized to-date: I-F3 (*3, 5*), I-B (*6*), and V-K CAST (*2*). One of the most remarkable aspects of CAST function is the precision of programmable insertions: DNA cargo is inserted in a single orientation and within a narrow window from the protospacer. These features appear to be generally conserved across CAST families (*3, 5–7*), and are likely a consequence of transpososome architecture (i.e. the nucleoprotein complex containing all CAST components).

Across all CAST elements (*4*), four or more proteins make up CAST transposition machinery: the CRISPR effector (Cas12k or Cascade), TniQ/TnsD, TnsC, and the transposase (TnsB or TnsA/TnsB). In all Tn7-like systems, a specialized CRISPR effector or DNA binding protein is hypothesized to recruit core transposition proteins sequentially (*8*). In CAST elements, TniQ protein associates with the target site via interactions with the CRISPR effector (*9*), and in some cases is shown to require a host factor(s) (*10, 11*). The AAA+ regulator TnsC serves as a ’matchmaker’ by interacting with both TniQ/TnsD and the transposase (TnsA/B) (*12, 13*). Very little structural information exists to explain how TnsC bridges the components in the hypothesized transpososome assembly. Furthermore, how core transposition machinery evolved to adopt different targeting strategies (*6, 14, 15*) remains an open question.

Due to its simplicity and robust *in vitro* activity (*2*), the V-K CAST system from *Scytonema hofmannii* (ShCAST) is an attractive system for understanding overarching mechanisms used by CAST elements. Though the structure of each ShCAST component has been individually characterized (*16–19*), crucial information on the spatial associations across the entire transposition recruitment process are missing (***Fig. 1A***). Using a DNA substrate representing the product of ShCAST integration, we reconstituted the ShCAST transpososome that was designed to mimic the natural configuration found with programmed integration (***Fig. 1B***). Using single-particle cryo-EM we obtained a high-resolution structure (3.5 Å), sufficient to accurately resolve interactions between all components within the nearly megadalton (961 kDa) nucleoprotein complex.

**Fig. 1.**
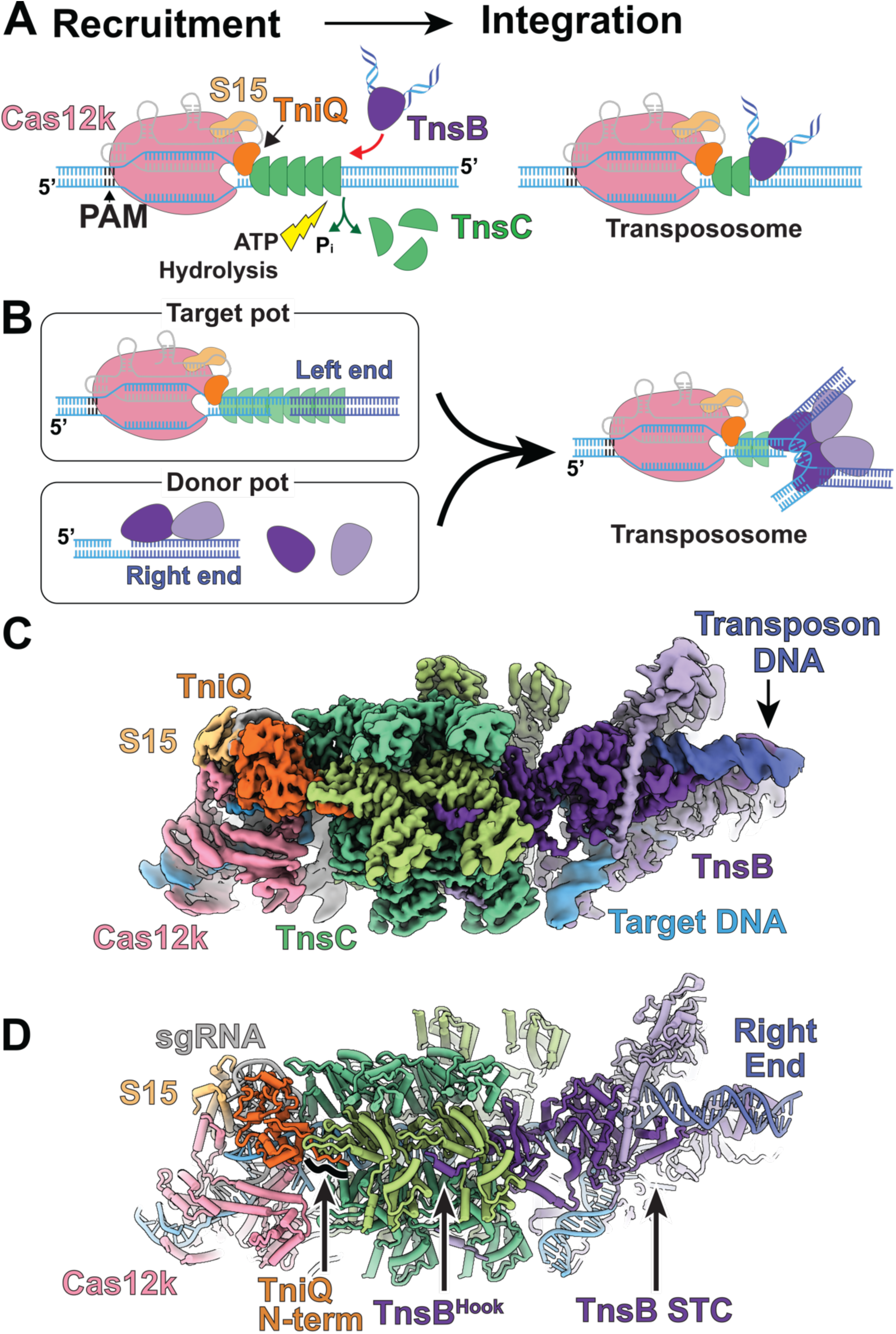
Cryo-EM structure of ShCAST transpososome. **A.** Mechanistic model of ShCAST recruitment (left) and integration (right). TnsC filaments (green semi-circle) associates with target site proteins: Cas12k (pink), TniQ (orange), and S15 (tan). The Protospacer Adjacent Motif (PAM, black) defines the beginning of the protospacer and R-loop, where the CRISPR effector, Cas12k and single guide RNA (sgRNA, shown in gray), binds. TnsB bound to transposon ends (purple) is recruited to the target site through interactions with TnsC (red arrow). ATP-hydrolysis (yellow lightning bolt) is stimulated by TnsB, resulting in the release of phosphate (labeled Pi, green arrow) and disassembly of TnsC (green arrow). Upon integration (right), a nucleoprotein complex consisting of all CAST components forms at the target site (right), referred to throughout as the transpososome **B.** Schematic for transpososome sample preparation. The target pot and the donor pot (left) were prepared independently to mimic the process of the RNA-guided transposition. DNA substrates for both the target pot and donor pot have target DNA region (light blue) and transposon-end region (dark blue) that are connected by 5 base pairs of single-stranded DNA, designed to form a strand-transfer complex (see Materials and Methods). Target pot and donor pot include target-site associated proteins (Cas12k, S15, TniQ, and TnsC) or TnsB, respectively, in addition to the corresponding DNA substrate. The transpososome (right) is reconstituted by combining the target and donor pot (see Materials and Methods). **C.** 3.5 Å resolution cryo-EM reconstruction of the ShCAST transpososome. Cryo-EM reconstruction is filtered according to local resolution. Each component in the complex is colored as defined in panel A. Different shades (light/dark green and light/dark purple) indicate different subunits of the same protein. Target DNA is colored light blue and transposon DNA is colored dark blue. **D.** Atomic model of the ShCAST transpososome. Features highlighted throughout the main text are labeled. The TniQ N-term corresponds to TniQ residues: 1 – 10. TnsB^Hook^ corresponds to residues: 570 – 584. STC is an abbreviation for strand-transfer complex (STC), see main text for details. Right end refers to the right end of the transposon DNA.

We discover that transpososome architecture promotes orientation-specific association across CAST components, in which TnsC has uniform polarity and dedicated faces of interaction with TniQ and TnsB, consistent with models of Tn7 transposition (*13, 20*). Cas12k, TniQ, and S15 stabilize R-loop formation, but TniQ and TnsC form the primary connection between CRISPR effector and transposase. We also observe novel TnsB-TnsC interactions, which are important for TnsB-promoted ATP hydrolysis. Importantly, in the context of the transpososome TnsC protomers make additional interactions with the target DNA distinct from that found in helical TnsC filaments(*19*) and are important for transposition activity. Finally, we discover that the transpososomes assemble with a slightly different number of TnsC protomers (+/− 1 protomer), hinting at the molecular basis of the intrinsic variability of ShCAST insertions. The structural insight gained from this transpososome structure contributes significantly to the understanding of CAST transposition and serves to identify avenues for protein engineering.

## Results

### DNA substrate design and transpososome reconstitution

ShCAST insertions occur within a 61-66 base pair window from the protospacer adjacent motif (PAM) in a single orientation (*2, 19*). We therefore designed a strand-transfer substrate (***Fig. S1***) that contains binding sites for all ShCAST components within this window (*2*). The ‘target pot’ from the transpososome reconstitution procedure(***Fig. 1B***) is similar to the procedure used to reconstitute a ShCAST subcomplex referred to as the ’recruitment complex’ (*10*), containing all components except TnsB. We also based our reconstitution procedure on established methods for monitoring transposition *in vitro* (*19, 21*) and reasoned that this procedure would mimic the recruitment and integration process (***Fig. 1B***). Incubation of target DNA with target site binding proteins (Cas12k, TniQ, S15, and TnsC) and ATP resulted in a heterogeneous assembly, consisting of different TnsC filament lengths (as assessed by negative stain EM, ***Fig. S2A***). Subsequent incubation with transposon-DNA bound TnsB, followed by enrichment for DNA-bound complexes resulted in assembly of complete transpososome particles (***Fig. 1B & S2B***), which were subjected to high-resolution cryo-EM imaging and image analysis (see Material and Methods for details, ***Fig. S3***).

### Transpososome architecture reveals orientation-specific associations across CAST components

A 3.5 Å Cryo-EM reconstruction of the reconstituted ShCAST transpososome reveals an assembly composed of one Cas12k, one TniQ subunit, one ribosomal S15 subunit, four subunits of TnsB and two full turns of TnsC (***Fig. 1C***). We were able to distinguish two different transpososome populations containing either 12 or 13 TnsC protomers (the former is shown in Fig. 1). Because particles containing 12 TnsC protomers were most frequently observed (73% of particles used in final refinement) and yielded a slightly higher resolution cryo-EM map (3.5 Å vs 3.7 Å), we refer to this configuration as the ’major’ configuration and focus on it throughout subsequent sections unless otherwise noted (***Fig. S3***).

The ShCAST recruitment complex is heterogeneous, consisting of both ’productive’ and ’nonproductive’ complexes, distinguished by the direction of TnsC filaments bound to DNA (*10*). Notably, the number of TnsC turns in the recruitment complex are also variable, consistent with our observations (***Fig. S2A***). In contrast to the heterogeneity of the recruitment complex, transpososome particle images reveal uniform TnsC polarity (***Fig. 1C-D***), in which the N-terminal face of TnsC interacts with TniQ at the target site and the C-terminal face of TnsC interacts with TnsB (***Fig. S4***). In addition, TnsC filaments in transpososome particles are of uniform length, consisting of two turns of TnsC. TnsC is disassembled by the transposase (TnsB) during recruitment (***Fig. 1A***) (*19*). Therefore, the homogeneous TnsC polarity and length observed in our particle images suggests that: 1. TnsC not forming productive interactions with target site proteins are presumably disassembled by TnsB, and 2. TnsB disassembles TnsC filaments until a certain point, reflected in the uniform length of TnsC filaments observed in our transpososome reconstitution. Therefore, the interactions between TnsC and target site associated proteins (Cas12k, TniQ, and S15) are stabilized against further disassembly by TnsB (***Fig. 1A, right***). Thus, TnsC’s matchmaking function is architecturally enforced in the transpososome by uniform TnsC polarity and the face-specific association of CAST components TniQ and TnsB.

### TniQ interactions with TnsC are critical for CAST transposition

Consistent with TniQ’s functional importance in ShCAST transposition (*17*), TniQ is located at the PAM-distal end of the R-loop close to Cas12k and S15 (***Fig. S5***) and appears to primarily interact with TnsC and RNA, consistent with the productive recruitment complex (*10*). We observe that TniQ interacts most extensively at the second TnsC protomer from Cas12k, TnsC11, but is always observed to bridge the two TnsC protomers closest to Cas12k (***Fig. 2A, S6***). More specifically, the N-terminal tail of TniQ forms structured interactions with TnsC11’s finger loop and is generally hydrophobic in nature (***Fig. 2B***). From the structure, we identified the following important residues: hydrophobic residues W10 and F12 which are buried into a hydrophobic cleft in TnsC11 formed by the finger loop (***Fig. 2B***). H34 forms hydrogen bonding interactions with acidic residue E131 on the N-terminal face of TnsC11 (***Fig. 2C***). Truncation of the first 10 residues or mutation of the hydrophobic residues (W10, F12, and H34) results in near complete loss of transposition activity (***Fig. 2D***), consistent with the essential role suggested by our structural observations. TniQ interactions with the terminal TnsC protomer (TnsC12) have a much less extensive interaction interface compared to TniQ-TnsC11 interactions (325 Å^2^ vs 915 Å^2^, see Materials and Methods). Consistent with the reported importance of S15 for formation of the productive recruitment complex (*10*), we find that S15 generally improves transposition activity. However, addition of S15 does not change our conclusions regarding the importance of TniQ residues (***Fig. 2D***). Cas12k-TnsC interactions are completely absent (***Fig. S7***), and 3D variability analysis suggests that Cas12k allows flexible linkage to the rest of the ShCAST components (***Movie S1***). Together, these observations collectively suggest that the TniQ-TnsC interactions we observe are the primary connections required for directing insertions to a guide RNA-directed target site. This supports the idea that engineering new associations with target-site recognition modules (i.e. Novel non-CAST CRISPR effectors) can be achieved by focusing on TniQ.

**Fig. 2.**
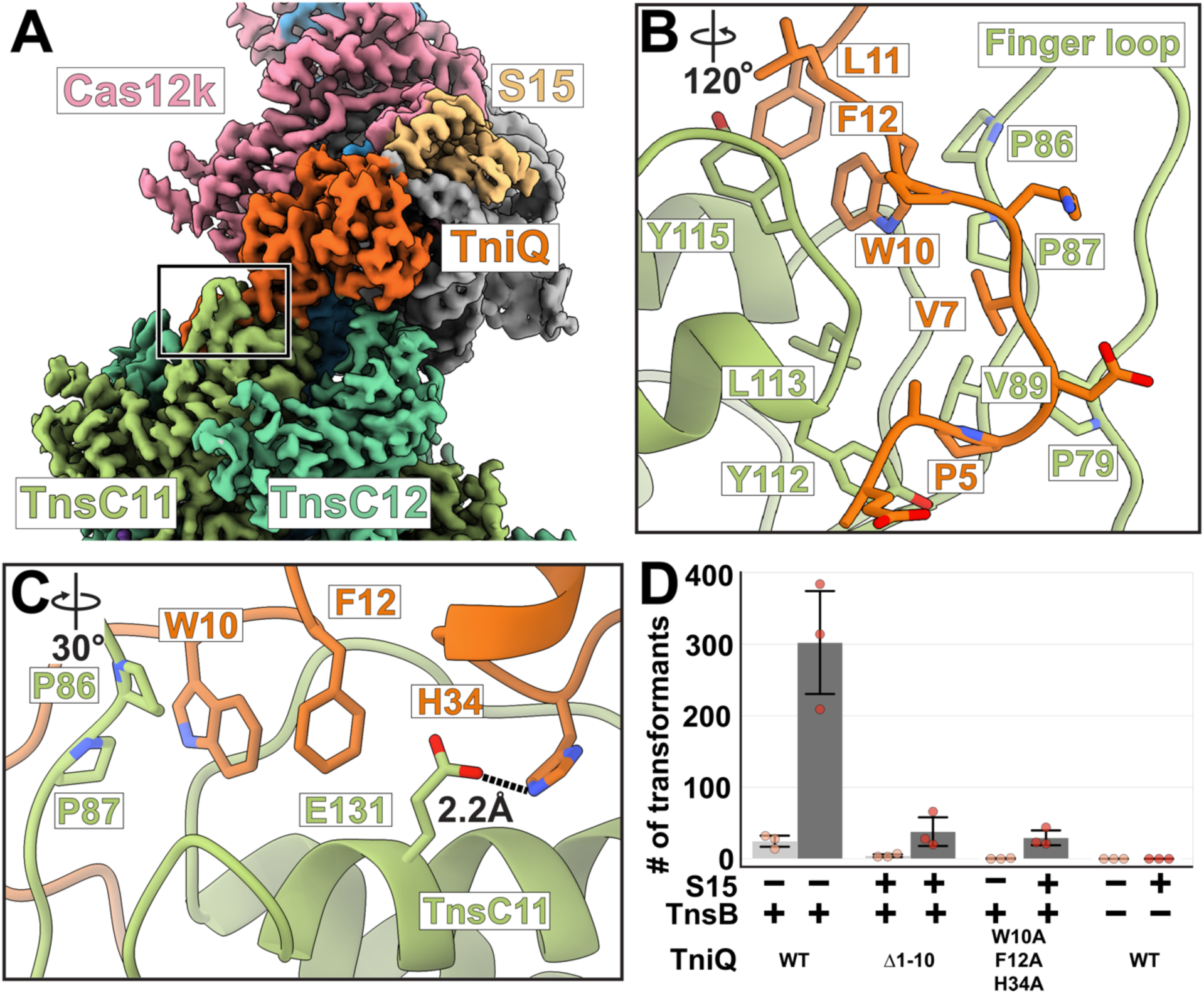
TniQ-TnsC interactions are crucial for ShCAST transposition. **A.** The composite cryo-EM map shown is created from two separate cryo-EM maps produced from local refinement (see Materials and Methods for details). The composite map is colored as previously described in Fig.1. TnsC protomers are labeled according to their position in the transpososome (TnsC1 is the closest TnsC protomer to TnsB). Black box indicates the TniQ-TnsC interface shown as insets in panel B and C. **B-C.** Atomic model of TniQ-TnsC interactions. Two different views are shown: rotation with respect to the inset in panel A is indicated at the top left of each panel. **B.** N-terminal tail of TniQ is shown (in sticks, with specific residues labeled) interacting with a hydrophobic cleft created by the finger loop of TnsC (labeled). **C.** Hydrogen bonding interactions are shown in dashed black lines along with distance (in Å) between the donor and acceptor atoms. **D.** TniQ mutations eliminate *in vitro* transposition activity, consistent with the interactions observed in our transpososome structure. Inclusion (dark gray) or exclusion (light gray) of S15 and TnsB is indicated with + or −, respectively. Wild-type TniQ is either indicated with WT or labeled according to the mutation made. Δ1-10 refers to TniQ construct missing the first ten residues of its N-terminus. The number of transformants is plotted for each condition tested as a proxy for the transposition activity. Data are represented by the mean; error bars indicate SD (n = 3, biological triplicates). Raw data points are shown in red.

### TnsB-TnsC interactions within the transpososome are precise and well-defined

We next focused on the interactions between TnsB and TnsC in the transpososome structure (***Fig. 3A***). TnsB is a RNaseH transposase that bears significant similarities to MuA from bacteriophage Mu (*18, 22*). TnsB contains multiple functionally distinct domains dedicated to DNA-binding, catalytic activity, or interactions with the AAA+ regulator TnsC (*18*) (***Fig. S8***). The previously determined tetrameric TnsB strand-transfer complex (STC) structure (PDB: 7SVW) (*18*) docks well into the transpososome cryo-EM map, requiring only minimal modifications throughout (1.54 Å Cα r.m.s.d. ***Fig. S9***). Full-length TnsB has a 52 residue long flexible linker that is predicted to be disordered (residues 518 - 569), that connects the STC to the C-terminal ’Hook’ (TnsB^Hook^, 570-584) (***Fig. S8***). TnsB^Hook^ serves as the point of contact with the TnsC filament body and is critically important for CAST transposition (*18*). We identified well-defined density corresponding to the TnsB ’Hook’ (TnsB^Hook^,) at four of the six TnsC protomers closest to TnsB (TnsC protomers 2-5, ***Fig. 3B & Fig. S10***). After docking in these structurally characterized portions of TnsB, we discovered additional unassigned density corresponding to TnsB residues 475 - 542, which forms a helix-turn-helix motif (***Fig. 3C***). Therefore, although TnsB-TnsC associations may be in principle flexible, we see very well defined TnsB-TnsC interactions in the transpososome, with all four TnsB protomers engaged at precise locations on the TnsB-proximal TnsC hexamer.

**Fig. 3.**
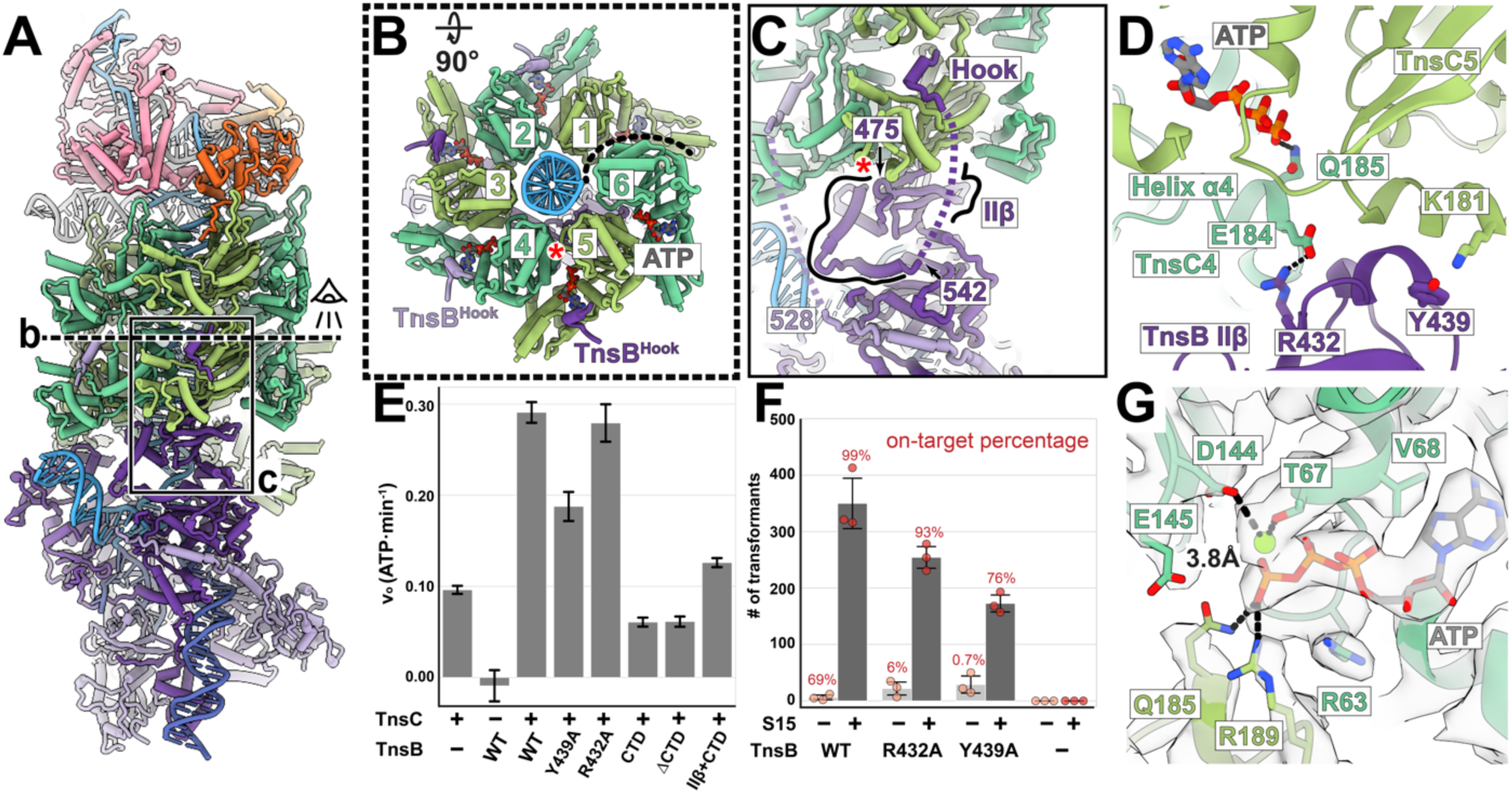
Transpososome structure reveals TnsB-TnsC interactions that are important for recognition of TnsC and stimulation of TnsC’s ATP hydrolysis activity. **A.** Overview of TnsB-TnsC interactions in the transpososome structure. Dashed line indicates the slice of the transpososome structure visualized in panel B, and the eye indicates the viewing direction. Box indicates the inset region shown in panel C. **B.** The N-terminal face of the TnsC hexamer closest to TnsB. TnsC protomers are labeled according to their position in the transpososome (index starts at 1 from the TnsC protomer closest to TnsB). Dashed line indicates where hexamer ends. Colors follow convention established in Fig. 1: DNA is shown in blue, TnsC protomers in green, and TnsB^Hook^ in purple. Four TnsB^Hook^ peptides bind at TnsC protomers 2 - 5. TnsB^Hook^ is colored according to the TnsB protomer it is closest to. Red asterisk (*) indicates where domain IIΔ (residues 410 – 474) associates across two TnsC protomers on the C-terminal face of TnsC. ATP is shown in stick representation in each of the ATP binding pockets between TnsC protomers. **C.** Additional side view showing details of TnsB-TnsC interactions. Residue positions 475 - 542 (labeled) correspond to the additional structured TnsB domain (highlighted with black lines) observed to interact between two TnsC protomers at the C-terminal face of the TnsC hexamer shown in panel B. Red asterisk (*) indicates the beginning of Helix α4 (described in further detail in D). Dotted lines indicate the flexible linker (not observed in our structure). Assignments are based on spatial proximity. **D.** TnsB-TnsC interactions near the ATP binding pocket. Domains and secondary structure elements referenced in the main text are labeled along with TnsC protomers. Possible interacting residues are shown in stick and labeled. Possible hydrogen bonds are shown as black dashed lines. **E.** Alterations to TnsB domain IIΔ decrease ATP hydrolysis activity, as assessed using an ATP hydrolysis assay (see Materials and Methods). Inclusion or exclusion of TnsC is indicated with + or −, respectively. Exclusion of TnsB is indicated with −, TnsB is either labeled WT (for wild-type) or labeled according to the mutation made. CTD refers to TnsB residues: 476 – 584. ΔCTD refers to TnsB construct missing CTD residues: 476-584. IIΔ+CTD construct refers to TnsB residues 410 – 584. Hydrolysis rate (vo, ATP·min^-1^) is shown for each variant tested (see Materials and Methods for details), and error bars indicate SD (n=3, biological triplicates). **F.** *In vitro* transposition assay testing TnsB mutants in the presence (dark gray) or absence (light gray) of S15. Presence or absence of each component is indicated with + or −, respectively. Wild-type TnsB is either indicated with WT or labeled according to the mutation made. Number of transformants is plotted for each condition tested as a proxy for the transposition activity. Data is represented by the mean; error bars indicate SD (n = 3, biological triplicates). Raw data points are shown in red. On-target percentage of the transposition was estimated from the Illumina sequencing and indicated on the corresponding bar plot (see Materials and Methods for details). **G.** TnsC within the transpososome is in the ATP-bound state. Cryo-EM density is shown in transparent gray surface. Two different TnsC protomers are shown, colored in light and dark green. Magnesium ion is shown as green sphere. ATP is shown in gray. Functionally important residues are labeled and shown in stick representation. Hydrogen bonds are indicated with dashed black lines. Distance from D144 to Magnesium is indicated in Å.

### TnsB interactions with TnsC’s C-terminal face are related to stimulation of ATP hydrolysis

In the ShCAST system, the transposase TnsB is recruited to the target site via TnsC (***Fig. 1A, left***) (*18*). This is intrinsically tied to ATP-hydrolysis, because substitution with a non-hydrolyzable ATP analog (AMP-PNP) inhibits TnsB-mediated filament disassembly and results in non-targeted transposition (*19*). Therefore, a crucial role of TnsB is also to stimulate TnsC’s ATP-hydrolysis activity in the process of target site selection. Two TnsB-TnsC interactions have been biochemically demonstrated to be required for this process, one of which corresponds to TnsB^Hook^ and the other one located elsewhere in full-length TnsB (*18*). Excitingly, within the transpososome structure we observe a second TnsB-TnsC interaction, not previously observed, at the C-terminal face of TnsC (***Fig. 3C-D***). In particular, residues 410 - 542, (including domain IIΔ) are positioned to bridge two TnsC protomers (***Fig. 3C***). Intriguingly, domain IIΔ localizes close to TnsC’s ATP-binding pocket, abutting against helix α4 in TnsC (***Fig. 3D***). This helix contains functionally important, conserved residues Q185 and R189 (***Fig. 3D***), which ’sense’ the nucleotide-bound state (*19*).

Although the features of this TnsB-TnsC interaction are suggestive, TnsB is not actively promoting ATP hydrolysis in the configuration observed here. To test the idea that the TnsB-TnsC interactions we observe in the transpososome structure are required for TnsB’s ability to stimulate TnsC ATP-hydrolysis (and trimming of the TnsC filament), we tested various mutation constructs using a malachite green ATP hydrolysis assay (***Fig. 3E***, see Materials and Methods). Consistent with expectations, TnsC has low basal levels of ATP-hydrolysis activity on its own, and wild-type TnsB stimulates TnsC’s ATP hydrolysis activity substantially (***Fig. 3E***). The Y439A TnsB mutation at the TnsB-TnsC interface (***Fig. 3D***) reduced TnsB’s ability to stimulate ATP-hydrolysis, but the other mutant (R432A, ***Fig. 3D-E***) had no effect. To test whether domain IIΔ is generally required, we next tested various TnsB fragments for their ability to stimulate ATP hydrolysis. TnsB C-terminal constructs (called CTD, residues 476-584) and TnsB C-terminal truncations (ΔCTD, residues 1-475) both fail to stimulate ATP hydrolysis, as expected based on our previous work (*18*). In contrast, C-terminal constructs including domain IIΔ (IIΔ+CTD, residues 410-584) increased ATP-hydrolysis levels over basal levels of TnsC ATPase activity (i.e. no TnsB) and CTD constructs (TnsB residues 476-584) (***Fig. 3E***). Despite these effects on ATP-hydrolysis, TnsB mutations at this interface only slightly reduced transposition activity compared to wild-type levels (***Fig. 3F***). However, consistent with the idea that these TnsB mutants have a reduced capacity to stimulate ATP hydrolysis, targeted transposition (as assessed by Illumina sequencing, see Materials and Methods) is significantly reduced in the absence of S15 (69% vs ≤ 6%, ***Fig. 3F & S11***). The addition of S15, a host factor shown to boost overall transposition activity (*10*), results in a less dramatic effect (99% vs 76-93% on site targeting percentage, ***Fig. 3F & S11***), consistent with the idea that S15 can generally promote targeted transposition by stabilizing CRISPR-effector binding. It is particularly striking that in the absence of S15, WT TnsB retains high levels of on-site targeting (69%) whereas mutant TnsB insertions are essentially random (***Fig. S11***). These dramatic effects suggest that the function of TnsB to promote target-site selection through TnsC filament disassembly is compromised, similar to the effects observed with non-hydrolyzable analog, AMP-PNP (*19*).

In this context, the defined assembly of TnsC we observe in the transpososome may correspond to either an ATP hydrolysis-resistant state or a stable post-hydrolysis state. To distinguish between these two possibilities, we used local refinement (focusing on TnsC) to generate a 3.2 Å map (***Fig. S12***), sufficient to unambiguously distinguish ATP and ADP. This improved map unequivocally shows that ATP is bound at all protomers (***Fig. 3G & Fig. S13***). Our observations are consistent with the idea that although TnsB stimulates ATP-hydrolysis and TnsC filament disassembly, aspects of the transpososome structure likely prevent hydrolysis, allowing the stable configurations observed here.

### TnsC structure in the transpososome deviates from previous helical TnsC structures

In the absence of additional factors, ATP-bound TnsC forms continuous helical filaments (referred to throughout as helical TnsC) with DNA, and has no preference for filament length (*16, 19*). However, the precise insertion profiles of ShCAST suggest that TnsB trims back TnsC filaments to a specific length prior to transposon DNA integration (*2*). Thus, we wondered whether significant structural differences distinguish helical TnsC from the structure observed here. We discover that the two full turns of TnsC present in the transpososome are overall highly similar to helical TnsC (overall Cα r.m.s.d. with PDB: 7M99 is 1.6 Å). However, not all of the TnsC protomers follow the helical symmetry of the ATP-bound TnsC filament (***Movie S2, Fig. S14***), and imposing helical symmetry during refinement results in lower resolution reconstructions with aberrant features (3.7 Å, ***Fig. S15***), consistent with the idea that the configuration of TnsC in the transpososome is not technically helically symmetric. The largest structural changes in the assembly correspond to the TnsB-proximal (TnsC1, ***Fig. 4A***) and Cas12k-proximal (TnsC12, ***Fig. 4A***) TnsC protomers, which are collapsed towards the center of the TnsC assembly with Cα r.m.s.d. deviations between 2.1-2.5 Å (***Fig. 4A***). This suggests that the slight conformational changes observed here are due to interactions with other CAST components within the transpososome, namely TniQ and TnsB.

**Fig. 4.**
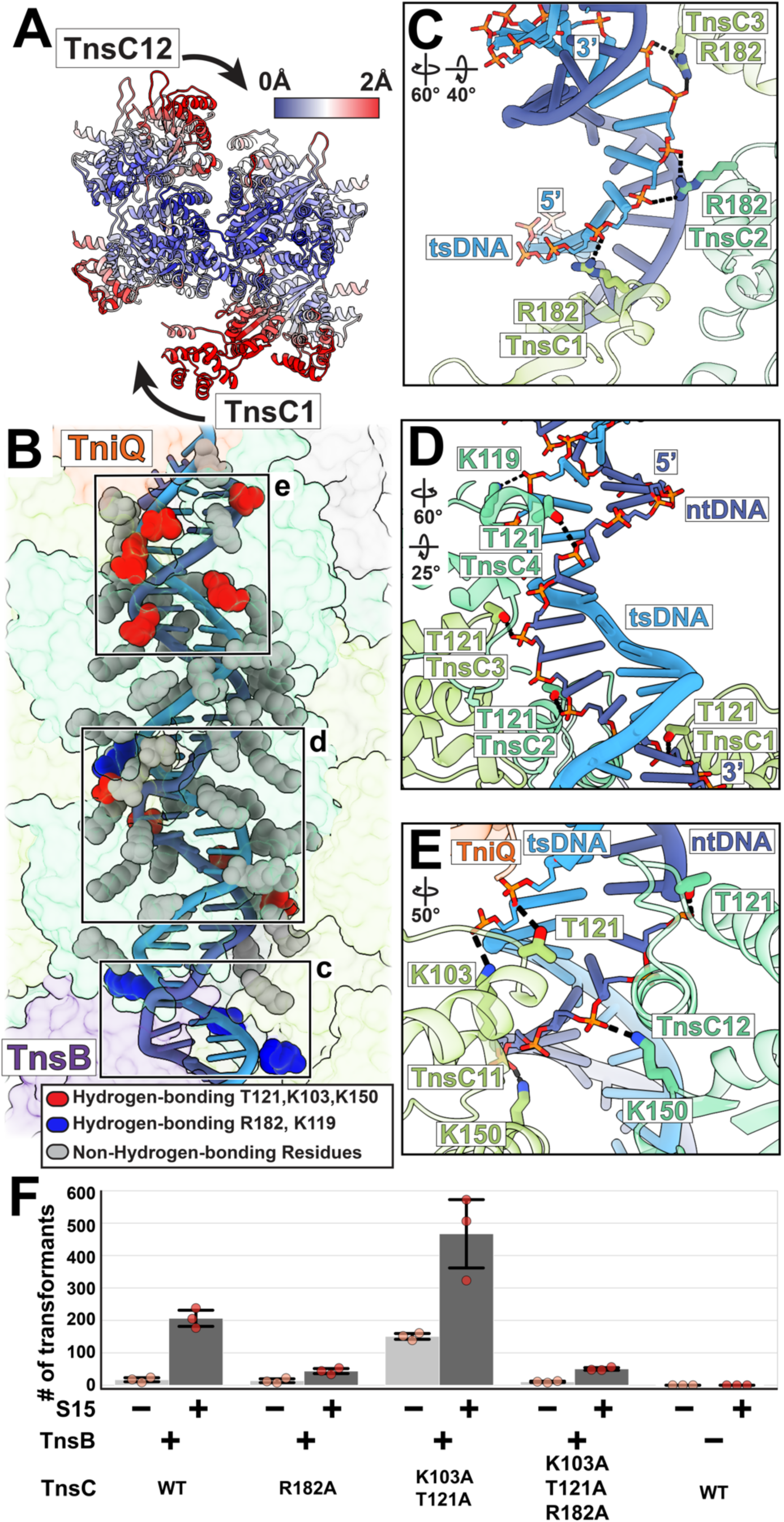
TnsC in the transpososome makes a network of important target DNA contacts throughout the transpososome. **A.** TnsC protomers in transpososome are colored by changes in Cα r.m.s.d. between the helical TnsC (PDB: 7M99) and TnsC in transpososome. Red indicates the largest changes (>2Å) and blue indicates small/no changes. Legend at the top right indicates colors assigned to r.m.s.d. in Å. The arrows represent the inward movement of TnsC, starting from the helical filament. **B.** TnsC-DNA interactions throughout the ShCAST transpososome. The TnsC protomers (green), TnsB (purple), TniQ (orange), and sgRNA (grey) are shown in transparent atomic surface. TnsC residues forming hydrogen-bonding interactions with DNA, ≤4Å away, are represented by spheres and colored either red (K103,T121, and K150) or blue (R182 and K119). Red residues are previously identified, functionally important residues that make novel contacts in the transpososome. Blue residues are newly identified residues that make functionally important, specific interactions with target DNA in the context of the transpososome. TnsC residues in close proximity to DNA but not forming specific interactions with the DNA backbone, >4Å away, are represented by spheres and colored grey. The target strand DNA (tsDNA, light blue), and non-target strand DNA (ntDNA, dark blue), are represented in ribbon. Black box indicates the TnsC-DNA interactions closest to Cas12k and TnsB shown as insets in panels C, D, and E. **C.** TnsB-proximal TnsC protomers (from TnsC1 to TnsC3) are shown interacting with the sugar-phosphate backbone of tsDNA (shown in stick). Hydrogen bonding interactions between protein residues (R182) and the sugar-phosphate backbone of DNA are represented with the dashed line. Rotations in top left show relationship of inset to overview in B. **D.** TnsB-proximal TnsC protomers (from TnsC1 to TnsC4) are shown interacting with the sugar-phosphate backbone of ntDNA (shown in stick). Hydrogen bonding interactions between protein residues (T121 and K119) and the sugar-phosphate backbone of DNA are represented with the dashed line. Rotations in top left show relationship of inset to overview in B. **E.** TnsC protomers (from TnsC11 to TnsC12) adjacent to Cas12k interact with both tsDNA and ntDNA immediately downstream of the R-loop. Hydrogen bonding interactions between protein residues (K103,T121, and K150) and the sugar-phosphate backbone of DNA are represented with the dashed line. Black dashed lines indicate hydrogen-bonding interactions. Rotations in top left show relationship of inset to overview in B. **F.** *In vitro* transposition assay of TnsC residues observed to interact with DNA. Inclusion or exclusion of TnsB is indicated with + or −, respectively. Inclusion of S15 (dark gray) or exclusion (light gray) changes overall activity levels but does not change the effect of mutant phenotypes. Wild-type TnsC is either indicated with WT or labeled according to the mutation made. Number of transformants is plotted for each condition tested. Data are represented by the mean; error bars indicate SD (n = 3, biological triplicates). Raw data points are shown in red.

### TnsC-DNA interactions differ across each of the two turns of TnsC in the transpososome

Given the lack of significant structural rearrangements of TnsC within the transpososome, we were especially interested to see that TnsC-DNA interactions appear significantly altered in the context of the transpososome compared to previous helical TnsC structures. In previous work, the helical symmetry of TnsC tracks with that of the bound DNA duplex (*16, 19*). In doing so, helical TnsC imposes its own helical symmetry, which distorts the bound DNA through underwinding (*16, 19*). In contrast to helical TnsC structures, within the transpososome TnsC does not track specifically with either strand of the DNA duplex (***Fig. 4B***). Comparing the structure of transpososome target DNA to B-form DNA, we discover that target DNA is only slightly underwound compared to B-form DNA (11 vs 10 base pairs per turn) and does not match the layer line spacing of TnsC in the transpososome (∼40 Å, ***Fig. S16***).

Due to the mismatch in TnsC (40 Å) and DNA (36 Å) repeat lengths, the specific TnsC-DNA interactions made in each of the two TnsC turns in the transpososome are distinct (***Fig. 4B***). In helical TnsC structures, residues K103 and T121 are consistently found to track with one particular strand of duplex DNA (5’ to 3’ in the direction of the C-terminal to N-terminal face, ***Fig. S17***) (*19*), which we confirm in improved, high-resolution (3.5 Å) cryo-EM reconstructions of TniQ-TnsC (***Fig. S18-19***). In the context of the transpososome, we now see a network of interactions, some of which correspond to new interactions by previously identified DNA-binding residues (T121, K103, and K150, red, ***Fig. 4B***) or new interactions made by residues not previously associating with DNA (K119 and R182, blue, ***Fig. 4B***). The rest of the interactions throughout the two turns of TnsC correspond to general electrostatic interactions, as these residues are too distant to form specific hydrogen-bonding interactions with the DNA backbone (gray residues, ***Fig. 4B & Table S3***). Notably, we notice that the two different sets of residues forming specific interactions with the DNA backbone are generally segregated spatially, either towards the Cas12k-proximal end (red residues, ***Fig. 4B***) or towards the TnsB-proximal end (blue residues, ***Fig. 4B***). In particular, the new interactions with the DNA backbone, involving R182 (***Fig. 4C***) and K119 (***Fig. 4D***) appear compelling and are supported by cryo-EM density (***Fig. S20***). Residues previously shown to interact with DNA (T121 and K103) in helical TnsC are now associating with both target and non-target strands of the DNA duplex (***Fig. 4E***).

Based on this, we hypothesized that TnsC residue R182 is critical for ShCAST transposition. Consistent with this, both single (R182A) and triple mutants (K103A/T121A/R182A) significantly decreased overall transposition activity (***Fig. 4F***), while double mutant K103/T121 had an overall increase in transposition activity (***Fig. 4F***), consistent with our previous results (*19*). Mutant phenotypes were consistent regardless of whether S15 is added, further emphasizing the importance of R182. Because these interactions are not observed in the TniQ-TnsC structure (***Fig. S19***), this is not solely due to the binding of TniQ but must be related to R-loop formation, consistent with other reports (*10*). We hypothesize that these network of interactions correspond to the distinct roles that TnsC residues play in transpososome formation and stabilization, which may explain why two turns of TnsC are observed in the transpososome assembly seen here.

### Different transpososome configurations are distinguished by TnsC-DNA interactions

Aligning both transpososome reconstructions (major and minor configurations) based on TnsB reveals an additional TnsC protomer (TnsC13) next to Cas12k in the minor configuration, which contains one extra TnsC protomer compared to the major configuration (with 12 TnsC protomers) (***Fig. 5A***). Correspondingly, Cas12k is rotated by approximately 60° in the minor configuration (solid surface, ***Fig. 5A***) compared to the major configuration (transparent surface, ***Fig. 5A***). Comparing the two configurations, the specific TnsC-DNA interactions in protomers adjacent to TnsB are virtually identical, since the same interactions are made at the same positions on the target DNA substrate (***Fig. S21***). At first glance, the TnsC-DNA contacts adjacent to Cas12k appear similar as well. However, these contacts are shifted over by one nucleotide in the minor configuration compared to the major configuration (***Fig. 5B***). That is, the last two TnsC protomers in both configurations interact in the same way with TniQ, resulting in a translational shift of TnsC-DNA interactions (***Fig. 5B-C***). This leads to slight variability in the number of base-pairs contacted by TniQ (5 bp vs 4 bp, major vs minor, ***Fig. 5C***) and TnsC (30 bp vs 31 bp, major vs minor, ***Fig. 5C***). Further, the slight structural variations described here also suggests that more than one transpososome complex is stable, which suggests a mechanism to explain slight variability (∼5 bp) in ShCAST insertion profiles(*2*).

**Fig. 5.**
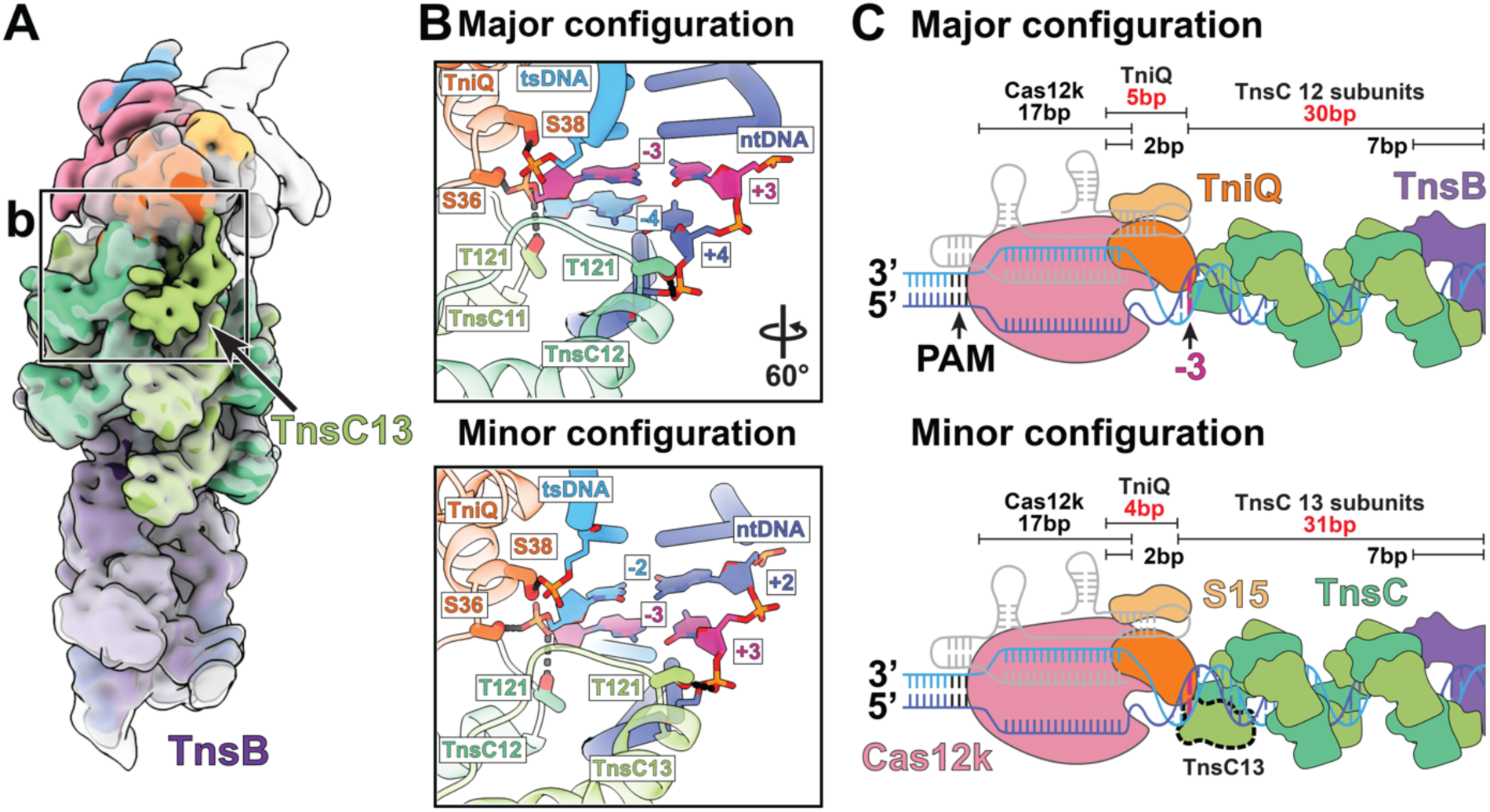
Two different transpososome configurations differ at TnsC’s N-terminal face. **A.** cryo-EM transpososome reconstructions, low pass filtered to 10Å, of both major and minor configurations aligned with respect to TnsB. The map corresponding to the minor configuration is colored and opaque while the major configuration is transparent. Black solid box indicates the TnsC-DNA interactions at the Cas12k face shown as inset in panel B. **B**. Major and minor configurations interact with both target -strand DNA (tsDNA) and non-target strand DNA (ntDNA). Residues from TniQ and TnsC in the minor configuration interact with the phosphate backbone that are one base-pair shifted towards R-loop compared to the major configuration. Hydrogen bonding interactions between protein residues and the sugar-phosphate backbone of DNA are represented with the dashed line. Base pairs interacting with the final two protomers of TnsC are represented in stick representation. Base pair positions at three bases downstream of R-loop (−3 of tsDNA and +3 of ntDNA) are colored magenta to serve as a landmark feature, facilitating comparison of the two configurations. **C.** Schematic of DNA binding contacts of each component (Cas12k, TniQ, and TnsC) in major (top) and minor (bottom) configurations. The number of base-pairs (bp) contacted by each component is listed and length indicated with double-bar lines. Positions that are interacting with two protein components (Cas12k and TniQ or TnsC and TnsB) are indicated as overlapping bars. Minor configuration includes additional TnsC protomer (TnsC13, dashed outline) proximal to Cas12k, which makes the DNA-binding contribution of TniQ and TnsC different in the two configurations (highlighted in red). The third base-pair downstream of the R-loop is colored magenta, identical to panel B.

## Discussion

Our structural observations allow us to fill in crucial mechanistic details concerning the recruitment and insertion of transposon DNA at a CRISPR-defined target site (***Fig. 6***). Once a productive complex is formed consisting of target site proteins Cas12k, S15, TniQ, and TnsC (***Fig. 6A-B***), TnsB-TnsC interactions stimulate ATP hydrolysis and TnsC filament disassembly. Our structure also suggests that the TnsB-TnsC contacts required for filament disassembly involve the two TnsB domains mentioned here: TnsB’s C-terminal Hook and domain IIΔ (***Fig. 6C***). ATP hydrolysis (and TnsC filament disassembly) continues until TnsB encounters the two TnsC turns closest to Cas12k. We hypothesize that two turns of TnsC are required for transpososome formation with the ShCAST element (***Fig. 6D***), a state resistant to TnsB-mediated ATP hydrolysis (***Fig. 6E***) and stabilized by new TnsC-DNA contacts, K103, T121, K119, and R182. These contacts appear to stabilize a TnsC configuration that is not fully engaged with DNA compared to the helical TnsC filaments formed outside the transpososome. We suggest that TnsC interactions with DNA via residue R182 are responsible for allowing the filament to be trimmed down to a precise length to allow programmed spacing between the PAM and insertion site.

**Fig. 6.**
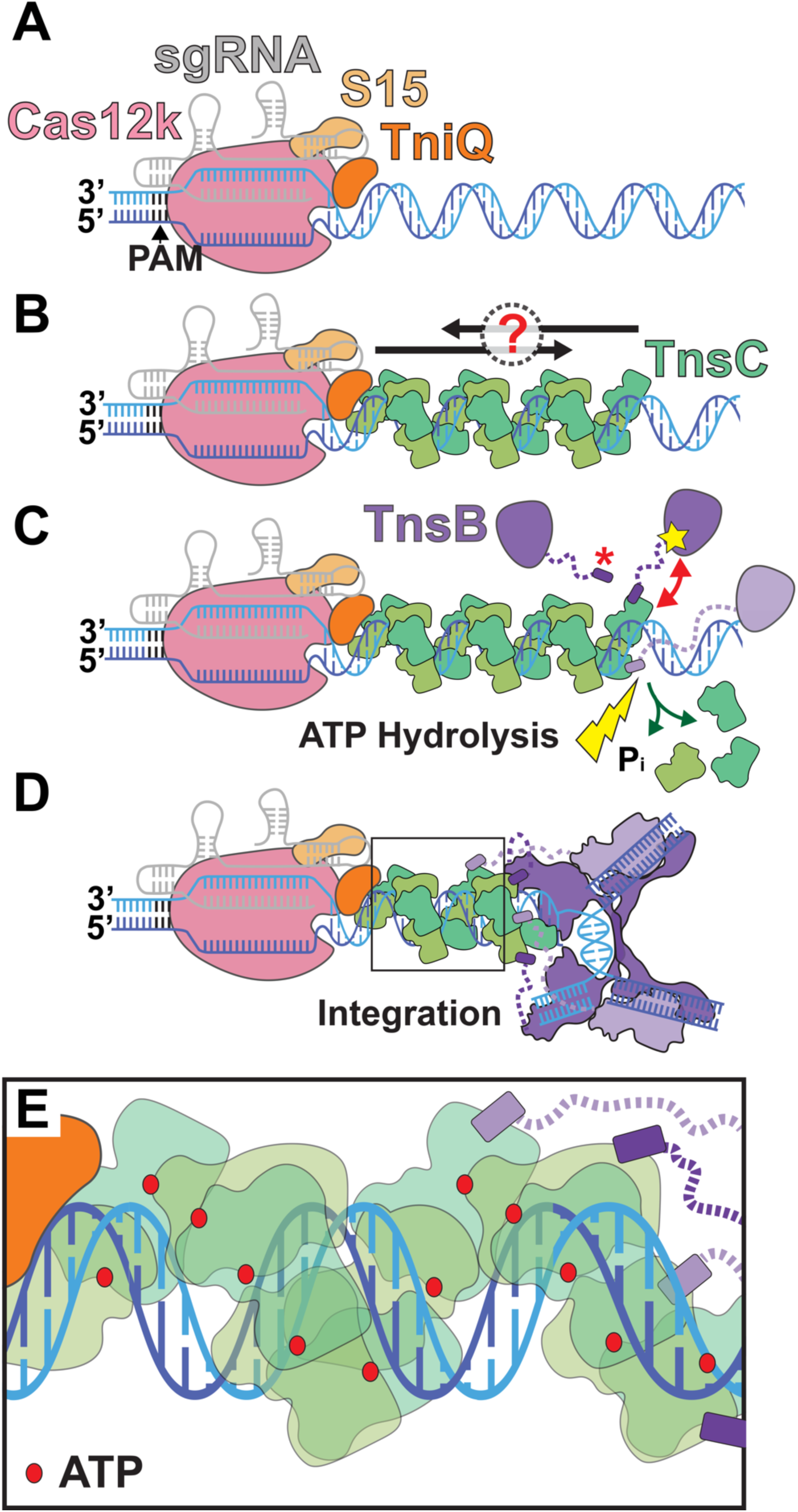
Mechanistic model of recruitment and integration of the transposition components in ShCAST. **A.** CRISPR effector Cas12k (pink) bound to sgRNA (gray) defines target site by forming an R-loop and associating with S15 (tan) and TniQ (orange). **B.** TnsC (green) may polymerize on DNA (blue) in two directions: towards or away from the target site (indicated by direction of the black arrow and question mark). TnsC interacts with target site associated proteins (Cas12k-sgRNA, S15, and TniQ) to form the recruitment complex. **C.** Two interactions between TnsB (purple) and TnsC: TnsB is recruited to TnsC filaments by TnsB^Hook^ (purple, indicated by red asterisk) and domain IIý (indicated approximately by yellow star) from TnsB can interact with TnsC (indicated by red double arrow), lead to disassembly of TnsC and promote ATP hydrolysis (yellow lightning bolt), resulting in the release of phosphate (labeled Pi, green arrows). Dashed line (purple) represents flexible linker between TnsB^Hook^ and the rest of TnsB. **D.** TnsB is recruited to the target site and forms strand-transfer complex (STC) upon integration. Four TnsB^Hook^ are bound to four TnsC protomers proximal to TnsB through the flexible linker (dashed purple line). Two turns of TnsC (12 protomers) are stably formed against disassembly. Together with target site associated proteins and TnsB STC, all CAST components form the transpososome at the integration site. **E.** A zoomed in view of the TnsC (boxed) in transpososome in panel D. All TnsC protomers are in ATP-bound (red circle) state. TnsC in the transpososome does not specifically track with DNA helical symmetry, unlike previous helical structures of TnsC.

This structure also reveals how ShCAST transpososome architecture conserves TnsC function across Tn7-like elements. ShCAST TnsC forms continuous helical filaments on DNA (*16, 19*), which are distinct from TnsC configurations found with prototypic Tn7 and I-F3 CAST subfamily. TnsC from prototypic Tn7 forms heptameric oligomers (*20*) and led to the original hypothesis that the ring would allow segregated interaction interfaces: the TnsD (a TniQ domain containing protein) interface maps to the N-terminal face, and the TnsA/B interface maps to the C-terminal face (*20*). Although I-F3 CAST TnsC also adopts a heptameric structure (*12*), it remains unknown whether TnsC in this system has similarly segregated interaction interfaces. However, ChIP-seq experiments suggest that TnsC plays a key role in associating with the Cascade complex, serving as a selectivity factor in subsequent recruitment of TnsA/B (*12*). Therefore, despite the structural differences in ShCAST TnsC, we hypothesize that the segregated interactions we observe in the ShCAST transpososome are conserved across Tn7-like transposition systems.

The segregation of interactions we observe in the transpososome also suggests a mechanism for how core transposition machinery can co-opt different CRISPR effectors via TniQ. This is particularly interesting in light of the I-B1 AvCAST elements, which contain features of both prototypic Tn7 and CAST elements (*6*). In particular, TnsC and the cognate transposase in I-B1 AvCAST elements can associate with two different TniQ domain proteins: TnsD (sequence-specific DNA-binding protein) or TniQ (associated with a CRISPR effector) in order to direct insertions (*6*). Competition between these two targeting modalities suggests that a single interface on TnsC is capable of interacting with either TniQ or TnsD (*6*). Our structure provides an elegant explanation for how this is achieved: by separating targeting machinery (TniQ/TnsD) from the core transposition machinery (TnsA/B) via segregated interactions on TnsC. This ensures that co-option of target-site binding proteins (i.e. with different CRISPR effectors or different DNA binding proteins) does not interfere with the function of core transposition machinery.

While the orientation and contacts we find in the transpososome assembly are remarkably uniform and well-resolved (***Fig. S2***) compared to the heterogeneity of other reconstitutions (*10*), we observe a slight variability in the oligomeric composition of TnsC within transpososome populations. This is not altogether unexpected, since CAST insertion profiles also contain a slight variability (∼5 base pairs) in their insertion profiles (*2*). Remarkably, the protein-protein associations we observe in either configuration are essentially identical, except TnsC and TniQ are shifted one nucleotide closer to Cas12k (***Fig. 5C***). Therefore, transpososome associations are constructed to allow minor flexibility. Taking this one step further, it is possible that the basis of CAST insertion variability is in the stable configurations of TnsC that preserve CAST component interactions. This suggests that more precise CAST elements can be engineered by reinforcing (i.e. stabilizing) specific spatial associations between CAST components within the transpososome.

## Acknowledgements

This work was performed at the National Center for CryoEM Access and Training (NCCAT) and the Simons Electron Microscopy Center located at the New York Structural Biology Center, supported by the NIH Common Fund Transformative High Resolution Cryo-Electron Microscopy program (U24 GM129539,) and by grants from the Simons Foundation (SF349247) and NY State Assembly. This work also relied on data collected using an instrument supported by the NIH through award S10OD030470-01. We additionally acknowledge XSEDE for computational resources used for image processing (MCB200090 to E.H.K). We acknowledge Shan-Chi Hsieh for sharing the protocol to analyze Illumina sequencing results. We thank Joseph Peters, Sam Sternberg, and Alba Guarné along with the Guarné, Peters, and Kellogg lab for feedback and stimulating discussion.

## Author Contributions

E.H.K. designed and supervised the project. J.U.P., A.W.T., and T.X.W. carried out protein expression and purification. R.D.S. cloned S15. J.U.P., A.W.T, and V.H.T prepared sample components and completed DNA-based pulldowns. J.U.P., A.W.T., V.T., A.N.R., and T.X.W. prepared GO grids and optimized the sample for cryo-EM imaging. J.U.P., A.W.T., and A.N.R. collected, processed, analyzed, and refined cryo-EM data. J.U.P. and A.N.R. built and refined atomic models. J.U.P. and A.W.T carried out all *in vitro* assays (ATP hydrolysis and *in vitro* transposition assays). J.U.P., A.W.T., A.N.R., and E.H.K. contributed to figures and manuscript writing. All authors contributed to manuscript writing and editing.

## Data and materials availability

Atomic models are available through the Protein Data Bank (PDB) with accession codes: 8EA3 (major configuration), 8EA4 (minor configuration) and 7SVU (TnsB^CTD^-TnsC-TniQ complex). All cryo-EM reconstructions are available through the EMDB with accession codes: EMD-27971 (major configuration), EMD-27972 (minor configuration), and EMD-25453 (TnsB^CTD^-TnsC-TniQ complex).

## Materials and Methods

### Protein cloning of ShCAST mutants and S15

TnsC (K103A, T121A, and R182A), TnsB (τιCTD; residues 1-475, CTD; residues 476-584, IIΔ+CTD; residues 410-584, R432A, and Y439A), and TniQ (τι1-10; residues 11-167, W10A, F12A, and H34A) mutants were cloned from the pXT130_TwinStrep-SUMO-ShTnsC vector (Addgene #135526), the pXT129_TwinStrep-SUMO-ShTnsB vector (Addgene #135525), and the pXT131_TwinStrep-SUMO-ShTniQ vector (Addgene #135527) respectively, using the Q5 site-directed mutagenesis kit (NEB) and two corresponding primers (forward and reverse primer pair are listed in **Table S2**). Double or triple mutants were generated by introducing mutation on top of the mutant plasmid vectors. For S15, the DNA sequence for *Escherichia coli* S15 was PCR amplified from *E. coli* K-12 MG1655 chromosomal DNA using the following primers: RPS15_Ext_Hifi_F and RPS15_Ext_Hifi_R (**Table S2**). This gene fragment was cloned into a linearized vector backbone (original ShTniQ vector from Addgene #135527) containing TwinStrep-SUMO at the N-terminus of S15 using the NEBuilder HiFi assembly protocol (NEB). TnsC clones were transformed into *Escherichia coli* BL21(DE3) competent cells (NEB), while TnsB and S15 clones were transformed into homemade T1 phage-resistant pRIPL BL21(DE3), derived from transforming pRIPL into BL21(DE3) Competent *E. coli* (NEB).

### Protein expression and purification of ShCAST components and S15

All clones except S15 were purified using previously described protocols (*1, 2*). For purification of S15, single colonies were inoculated in 10 mL of LB media containing 34 μg/mL chloramphenicol and 100 μg/mL ampicillin, and grown overnight at 37°C as starter cultures. The overnight starter culture was added to 1.5 L of TB medium containing the same ratio of antibiotics and grown until an OD of 0.4 at 37°C. The temperature was lowered to 20°C and the culture was induced at an OD of 0.6 with 0.4 mM isopropyl-β-D-thiogalactopyranoside (IPTG) and continued to grow overnight (16-18 hours). Cells were harvested by centrifugation at 5000 rpm for 10 minutes and resuspended in lysis buffer (50 mM Tris pH 7.4, 500 mM NaCl, 5% glycerol, 1 mM DTT, 1 mM PMSF, 0.1% NP-40, and protease inhibitors (Pierce, ThermoScientific)). The cell lysate was sonicated for 15 minutes with a pulse on for 2 s with 10 s rest in between. The sonicated lysate was centrifuged for 30 minutes at 10,000 rpm at 4 °C. The supernatant was loaded onto a strep-tactin superflow resin column (Qiagen) equilibrated with loading buffer (50 mM Tris pH 7.4, 500 mM NaCl, 1mM DTT, and 5% glycerol). The column was washed with 5 column volumes (∼15 mL) of wash buffer (50 mM Tris pH 7.4, 500 mM NaCl, 5% glycerol) before eluting with elution buffer (50 mM Tris pH 7.4, 500 mM NaCl, 5% glycerol and 4 mM desthiobiotin (SigmaAldrich)). The consolidated eluate was then incubated overnight at 4°C with SUMO protease (1/100 mass ratio of SUMO protease to S15 protein).

S15 was next purified using a heparin purification step. Protein from the strep-tactin elution (in 500mM NaCl) was diluted to a final salt concentration of 200 mM NaCl before loading onto a 1 mL heparin column (GE Healthcare) equilibrated in low-salt buffer (50 mM Tris pH 7.4, 200 mM NaCl, 1 mM DTT and 5% glycerol). The heparin column was washed with 20 column volumes of low-salt buffer, followed by a gradient elution from 200mM NaCl to 1M NaCl using high-salt buffer (50 mM Tris pH 7.4, 1.2M NaCl, 1 mM DTT, and 5% glycerol). Fractions containing S15 were concentrated, and buffer exchanged before size-exclusion chromatography was performed using a Superdex 200 increase 10/300 (Cytiva) equilibrated in the following buffer: 20 mM HEPES pH 7.5, 250 mM KCl, 1 mM DTT, and 5% glycerol. Fractions containing S15 were consolidated and concentrated to ∼2 mg/mL using a 6 mL 3K molecular weight cutoff centrifugal concentrator (ThermoFisher), before being aliquoted and flash-frozen in liquid nitrogen for long-term storage at -80°C.

### Preparation of DNA substrate

The general idea of the substrate design for the strand-transfer complex was described previously (*3*). However, instead of a symmetric strand-transfer DNA, we designed an asymmetric DNA that mimics the Cas12k-guided transposition product (***Fig. S1***). This design includes two DNA substrates: (i) target-pot DNA containing left-end (LE) sequence and protospacer and (ii) donor-pot DNA containing right-end (RE) sequence. Each DNA substrate was prepared independently. First, target-pot DNA substrate was prepared by annealing four synthetic oligonucleotides (IDT) of the following: target-LE_F, non-target_R, LE_R, and 5’ dethiobiotinylated LUEGO (*4*) (***Table S2***). These four oligonucleotides were mixed in a molar ratio of 1: 1.1: 1.1: 1. Second, donor-pot DNA is composed of three synthetic oligonucleotides including: RE_F, RE_R1 and RE_R2 (***Table S2***). These oligonucleotides were mixed in a 1: 1.1: 1.1 molar ratio. The mixture of oligonucleotides was supplemented with a 10X concentrated annealing buffer to make the final buffer composition the following: 10 mM Tris pH 7.5, 50 mM NaCl, and 1 mM EDTA. The mixture was then heated up to 95°C for 5 minutes and cooled down to 30°C at the rate of 1°C per minute using a thermal cycler (BioRad).

### Preparation of sgRNA

Using two primers, sgRNA_For and sgRNA_PSP1_Rev (***Table S2***), the DNA sequence encoding the T7 RNA polymerase promoter and sgRNA was amplified by PCR from pHelper_ShCAST_sgRNA vector (Addgene, #127921). The DNA template was then subjected to GeneJet PCR purification (ThermoScientific). sgRNA was produced by in vitro transcription using the HiScribe T7 High Yield RNA synthesis kit (NEB) with the PCR amplified DNA sequence as the template. Purified sgRNA was aliquoted and stored at -20°C.

### Reconstitution of transpososome complex and recruitment complex

The transpososome complex and recruitment complex were generated by stepwise assembly using a pull-down assay on streptavidin beads (Fig. 1B). For the target pot, annealed target-pot DNA was first bound to Streptavidin Mag Sepharose magnetic beads (Cytiva) equilibrated with reaction buffer (20 mM HEPES pH 7.5, 250 mM KCl, 15 mM MgCl_2_, and 1 mM DTT) and incubated at room temperature with shaking for 30 minutes. Cas12k and sgRNA were reconstituted in a 1:2 molar ratio and incubated for 30 minutes at 37°C in the following buffer: 20 mM HEPES pH 7.5, 160 mM NaCl, 15 mM MgCl_2_, and 1 mM DTT. The Cas12k-sgRNA complex was added to the target-pot DNA-bound beads and incubated for 30 minutes at 37°C in the presence of a 5-fold molar excess of S15 to DNA. The beads were washed three times with 500 μL wash buffer (20 mM HEPES pH 7.5, 250 mM KCl, 15 mM MgCl_2_, 0.05% Tween20, and 1 mM DTT) to remove excess protein and nucleic acids. 20-fold molar excess of TniQ and 10-fold molar excess of S15 to DNA were added to the beads and incubated at 37°C for 30 minutes. 10-fold molar excess of TnsC was diluted to a final salt concentration of 250 mM NaCl and added to the TniQ-S15 beads solution containing a final concentration of 1 mM ATP. The sample was incubated for 30 minutes at 37°C before the beads were washed three times with 500 μL wash buffer containing 1 mM ATP. The recruitment complex was prepared by eluting the target-pot DNA at this point using 50 μL of elution buffer (20 mM HEPES, 250 mM KCl, 15 mM MgCl_2_, 0.05% Tween20, 10 mM biotin, 1 mM ATP, and 1 mM DTT). For transpososome assembly, donor pot containing TnsB and donor-pot DNA was prepared independently in 6:1 molar ratio with the following final buffer composition: 25 mM HEPES pH 7.5, 100 mM NaCl, 15 mM MgCl_2_, and 1 mM DTT; and incubated at 37°C for 30 minutes. The reconstituted donor pot was added to the beads and incubated for 40 minutes at 37°C. Three washes of 500 μL wash buffer containing 1 mM ATP were performed before eluting the transpososome complex from the beads with 50 μL elution buffer. The eluate was diluted with the wash buffer containing 1 mM ATP for cryo-EM or negative-staining EM sample preparation by 6-10 fold or 15 fold respectively.

### *In vitro* transposition assay and high-throughput mapping of transposition events

All purified proteins (except Cas12k) were diluted to 2.5 µM using stock buffer (500 mM NaCl, 50 mM Tris pH 7.4, 10% glycerol, 0.5 mM EDTA, 1 mM DTT). Cas12k was assembled with sgRNA before incubation with other components in the following manner: sgRNA was added to Cas12k to make 2.5 µM Cas12k and 30 µM sgRNA. Separate reactions were prepared for the ‘target pot’ and ‘donor pot’. The ‘target pot’ contains 2.24 nM pTarget_pBC_KS+_PSP1 (a gift from the Peters Lab), 104 nM of Cas12k, 104 nM TnsC, 104 nM TniQ, and 1.25 µM sgRNA in the following transposition buffer (26 mM HEPES, 50 mM KCl, 0.2 mM MgCl_2_) supplemented with 2 mM DTT, 50 µg/mL BSA and 2 mM ATP. The ‘donor pot’ contains 1.08 nM pDonor_ShCAST_kanR (Addgene #127924), 104 nM TnsB, 2 mM DTT, 50 µg/mL BSA and 2 mM ATP in transposition buffer. Both pots were incubated at 37°C for 30 minutes before combining and adding MgOAc2 to a final concentration of 15 mM. The combined reaction was further incubated at 37°C for 2 hours. After the final incubation step, 20 μL of the reaction mixture was digested by 1 μL of Proteinase K (Ambion) and incubated at 37°C for 1 hour. 10 μL of the reaction was then transformed into 100 μL of Stellar^TM^ competent cells (Takara Bio USA) and plated on kanamycin plates. All the experiments were done in biological triplicates.

From the plates of each condition, the colonies were washed up and diluted to OD of 0.5 using LB. The liquid culture was then grown for 3 hours at 37°C before the plasmid extraction using a miniprep kit (QIAGEN). Prepared DNA was sequenced by the Microbial Genome Sequencing Center (https://www.seqcenter.com/) using the Illumina DNA sequencing service on the NextSeq 2000 platform. Paired-end reads (2×151bp) were analyzed using BBtools (http://sourceforge.net/projects/bbmap/). First, the reads were processed by BBDuk (BBtools) to collect adjacent target-DNA sequences from all the reads containing 30bp of the ShCAST left-end sequence. Then these sequences were mapped onto the pTarget_pBC_KS+_PSP1 using BBMap (BBtools) with a 90% sequence identity cutoff. Number of base pairs between the end of the PAM and the beginning of the ShCAST left-end sequence was used to define the position of the transposition. Based on the mapping, the transpositions at 55 – 70 base pairs downstream of the PAM were considered to be on-target transposition. The number of on-target transposition reads was divided by the total number of transposition reads to estimate the on-target ratio.

### Malachite green ATP hydrolysis assay

100 mL of malachite green reagent was prepared by the following protocol. First, 34 mg of malachite green carbinol base (ChemCruz) was dissolved into 40 mL of 1N HCl. 1 g of Ammonium Molybdate (MacronChemicals) was dissolved in a separate 14 mL of 1N HCl. These two solutions were mixed and diluted up to 100 mL using ddH_2_O. The reagent was then filtered using 0.45 μm syringe filter (VWR), and covered with an aluminum foil to avoid light. ATPase assay was completed using purified protein stocks of TnsC, and constructs of TnsB (Wild-type, ΔCTD, CTD, IIΔ+CTD, R432A, and Y439A). To minimize the processing time, 2X protein pot and 2X ATP pot were prepared independently. Buffer composition for both pots was the same as follows: 25 mM HEPES pH 7.5, 150 mM NaCl, 1mM DTT. In addition to the described buffer, 2X protein pot contains 10 μM TnsC, 2 μM TnsB, and 2.5 μM 60 bp DNA (annealed using 60bp_top and 60bp_bot, ***Table S2***), and 2X ATP pot contains 1 mM ATP and 2 mM MgCl_2_. 100 μL of each pot was mixed to initiate the ATP hydrolysis reaction. For each time point (10, 20, 30, 40, 50, 60, 90 and 120 minutes), 20 μL of the reaction mix was transferred to the 96-well plate (Corning) that contains 5 μL of 0.5 M EDTA to quench further hydrolysis. After taking all the time points, 150 μL of the room-temperature Malachite Green reagent was added to each well, and the plate was shaken for ∼5 minutes to develop the color for imaging. Absorbance at 650 nm was read from each well using an Infinite 200 plate reader (TECAN). For calibration, KH_2_PO_4_ solutions of the following concentrations were used to generate a calibration curve: 0 μM, 4 μM, 8 μM, 12 μM, 16 μM, 24 μM, 40 μM, and 60 μM. The data was plotted with time as the X-axis and the concentration of released inorganic phosphate as the Y-axis. The slope was obtained from the linear regression to get the reported vo (ATP·min^-1^) values. Three independent experiments were done for each condition, to generate the reported bar plot.

### Cryo-EM sample preparation and freezing of transpososome complex

The transpososome complex sample was prepared by diluting the elution from the pull-down by 6-, 8-, and 10-fold using wash buffer (described above) containing 1 mM ATP. Graphene oxide-coated grids were prepared following the protocol described previously (*3, 5, 6*). 4 µL of transpososome complex sample was loaded on the carbon side of a graphene oxide coated grid and incubated for 20 seconds in the Mark IV vitrobot chamber (ThermoFisher), which was set to 4 °C and 100% humidity. Each grid was blotted for 6 seconds with a blot force of 5, and then plunged into liquid ethane cooled with liquid nitrogen.

### Cryo-EM imaging and image processing of transpososome complex

Vitrified samples of transpososome complex were first screened using Talos Arctica (ThermoFisher) operated at 200 kV prior to large-scale data collection. Screened grids were imaged using a Titan Krios (300 kV, ThermoFisher) equipped with a BioQuantum energy filter (Gatan) and K3 direct electron detector (Gatan). 15,740 micrographs were collected using Leginon (*7*) at the 81,000X magnification (1.067 Å/pixel) using image shift, with the nominal defocus from -0.8 μm to -2.5 μm. Each movie was collected with 2 seconds of exposure with a total dose of 49.91 electrons/Å^2^, fractionated into 50 frames. Frames were aligned using MotionCor2 (*8*) through Appion (*9*), which was then imported to cryoSPARC (*10*) for CTF estimation and downstream image analysis. The workflow described below is shown in ***Fig. S3***. Using a template picker from cryoSPARC, initial particle stack was subjected to 2D classification in cryoSPARC. 2D classification resulted in 536,450 particles from selected 2D averages, which was subjected to *ab-initio* reconstruction in cryoSPARC. *Ab-initio* reconstruction separated particles into two classes, one containing 53% (285,090 particles) and another containing 47% (250,842 particles). The two classes were then subjected to homogenous refinement in cryoSPARC respectively. The particle stacks from both classes were subjected to 3D classification in RELION (*11, 12*), which removed junk particles with weak densities of Cas12k or TnsB. Each classification resulted in a particle stack that had well-defined configurations of ShCAST transpososome. The final particle stacks (188,055 particles for major configuration of transpososome and 67,096 particles for minor configuration of transpososome) were subjected to iterative CTF refinements (*13*) and Bayesian polishing (*14*) to improve resolution. The final maps of both major and minor configurations were then subjected to local refinement in cryoSPARC. Volume maps of Cas12k-S15-TniQ only, TnsC only, and TnsB only were generated in UCSF Chimera (*15*) using the volume eraser tool to remove the map. A mask for Cas12k-S15-TniQ, TnsC, and TnsB, respectively, was then generated in RELION (*11*) using the volume produced by Chimera as the input. Local refinement was then done in cryoSPARC (*10*) with the mask generated by RELION and the full map without particle subtraction. Local refinement of the major configuration of the transpososome resulted in 3.1 Å for the Cas12k-S15-TniQ region, 3.2 Å for both the TnsC and TnsB region. Local refinement of the minor configuration of the transpososome resulted in 3.6 Å for the Cas12k-S15-TniQ region, 3.6 Å for TnsC, and 3.8 Å for the TnsB region.

For Fig. 2, the composite map was generated using UCSF Chimera (*15*) command “vop maximum” with the aligned reconstructions from local refinements of Cas12k and TnsC. 3D Variability in cryoSPARC (*16*) was performed on the major configuration of the transpososome using the mask and particles from the final refinement. The filter resolution was set to 6 Å and the number of modes to solve was set to 3. After the job was completed, the 3D Variability display job using the particles and volumes from the 3D Variability job was completed with the output mode set to simple with 20 frames. The first series was then visualized in UCSF Chimera (*15*) using the “vop morph” command with all 20 frames. The results are presented in ***Movie S1***.

To impose helical symmetry on the two turns of TnsC in the transpososome, a mask was first created using a map of TnsC only (generated by the same protocol described above) as input in cryoSPARC (*10*). Helical refinement was then done in cryoSPARC (*10*) with the mask and the full map (major configuration) without particle subtraction. Initial helical parameters of the ATPγS-bound TnsC filaments (PDB: 7M99, rise = 6.82 Å and twist = 60°) (*2*) were used. Helical parameters were refined with the following range: 6.14 Å to 7.50 Å for the helical rise, and from 57° to 63° for the twist.

### Model building of transpososome complex

Initial models from previous studies, including PDB: TnsC (PDB: 7M99) (*2*), TnsB (PDB: 7SVW) (*3*) and sgRNA (PDB: 7PLA) (*17*), were first docked into the cryo-EM density and manually rebuilt using coot (*18*). Additional models were created using AlphaFold2 (*19*) for the following components: Cas12k, TniQ and S15, and rigid-body docked into the density. Coot was used to manually remodel or rebuild sections of the model. Real space refinement of the DNA substrate was completed in Phenix (*20*) with both base-pair and secondary structure restraints enforced. For measuring the interface area between TniQ and TnsC protomers, UCSF ChimeraX (*21*) command “measure buriedarea” was used with the desired two chains as inputs.

### Model Validation of transpososome complex

Map-model Fourier Shell Correlation (FSC) was computed using Mtriage in Phenix (*20*). Map-model FSC resolution of each dataset was estimated from Mtriage FSC curve, using 0.5 cutoff. The masked cross-correlation (CCmask) from Mtriage was reported as representative model-map cross-correlation. Model geometry was validated using the MolProbity (*22*). All the deposited models were submitted to MolProbity server to check the clashes between atoms, Ramachandran-plot, bond-angles, bond-lengths, sidechain rotamers, CaBLAM and C-beta outliers. All the model validation stats are summarized in **Table S1**.

### Helical parameter estimation using Rosetta

Helical parameters are estimated between the adjacent TnsC protomers within the complexes of the following: ATPγS-bound TnsC helical filament (PDB: 7M99), major configuration, and minor configuration of the transpososome. For example, from the major configuration of transpososome, helical parameters were estimated in between TnsC1 and TnsC2, between TnsC2 and TnsC3, between TnsC3 and TnsC4, and so on. Rosetta tool “make_symmdef_file.pl” was used for each pair of the TnsC protomers, with an example command presented at the bottom of this section. Obtained helical rises and turns were averaged to plot a bar-graph presented in ***Fig. S14***.

Rosetta command we used for helical parameter estimation is below: Rosetta/main/source/src/apps/public/symmetry/make_symmdef_file.pl -m HELIX -p input.pdb -a A -b B

### Reconstitution of TnsB^CTD^-TnsC-TniQ complex and imaging for cryo-EM

The C-terminal 109 residues from wild-type TnsB (termed hereafter as TnsB^CTD^) was cloned from pXT129_TwinStrep-SUMO-ShTnsB vector (Addgene, #135525) using Q5 site-directed mutagenesis kit (NEB) and two primers of following: TnsB_CTD_For and Ndel_primer_Rev (***Table S2***). Cloned TnsB^CTD^ was transformed to *E.Coli* BL21-RIPL competent cells (Agilent) and purified following the protocol for TniQ purification. DNA substrate was prepared by annealing two synthetic oligonucleotides of BCQ_top and BCQ_bot (***Table S2***). TniQ and TnsC were buffer exchanged into 25 mM HEPES pH 7.5, 200 mM NaCl, 2% glycerol, and 1 mM DTT (reaction buffer) prior to reconstitution of the complex. TnsC filament was first reconstituted by supplementing the reaction buffer with the following components: 128 μM TnsC, 8 μM DNA, 2 mM ATP, and 2 mM MgCl_2_. After 5 minutes of incubation on ice, 6-fold molar excess of TniQ was added to TnsC filaments and incubated at 37°C for 1 hour. TnsB^CTD^ was then added to the TniQ-TnsC mixture (4:1 molar ratio of TnsB^CTD^ to TnsC), followed by incubation at 37°C for 40 minutes. 4 μL of the TnsB^CTD^-TnsC-TniQ sample was loaded on UltrAuFoil R1.2/1.3 gold grids (Quantifoil) and vitrified using the Mark IV Vitrobot (ThermoFisher) set to 100% humidity and 4°C. Samples were blotted for 7 seconds with blot force 5, and then plunged into liquid ethane cooled with liquid nitrogen. Vitrified grids were imaged using Talos Arctica (ThermoFisher, 200 kV) with K3 direct detector (Gatan) and BioQuantum energy filter (Gatan), at 63,000X nominal magnification (1.33Å/pixel). SerialEM(*23*) was used for data acquisition with 3 x 3 image shift, and nominal defocus range from -1 μm to -2.5 μm. Total dose was set to 50 e^-^/Å^2^, which was fractionated into 50 frames.

### Image processing and model building for TnsB^CTD^-TnsC-TniQ complex

Beam-induced motion correction, CTF estimation, and initial particle picking were done using Warp (*24*) for the collected 1,271 movies. Initial particle stack from Warp was subjected to 2D classification cryoSPARC (*10*), to get subset of TniQ-bound particles to train topaz neural network (*25*), which resulted in the 795,631 particles. 2D classification resulted in 624,597 particles from selected 2D averages with high-resolution features, which were subjected to heterogeneous refinement in cryoSPARC (*10*). One class with better resolved TniQ (34%, 214,291 particles) was selected for downstream non-uniform refinement in cryoSPARC (*10*). The particle stack was subjected to 3D classification in RELION (*11, 12*) resulting in the intermediate particle stack with a stronger density of TniQ (70%, 150,358 particles). To improve the resolution of TniQ, this particle stack was subjected to two rounds of focused 3D classification (skipping alignments, tau fudge factor of 16), iterative CTF refinements (*13*) and bayesian polishing (*14*). This resulted in the final dataset of 61,515 particles, with the gold-standard FSC resolution of 3.5 Å. Due to the local variation of the map quality, LocSpiral (*26*) was used to post-process the map through COSMIC^2^(*27*). For model building of TnsC and TniQ, previously published structure of TnsC (PDB: 7M99)(*2*), and AlphaFold2 (*19*) generated TniQ were manually docked into the density, followed by manual editing using coot (*18*) and relaxation using Rosetta relax (*28*).

**Movie S1. 3D variability analysis visualizes a flexible association of Cas12k with respect to the rest of the transpososome.** Components in the transpososome were colored following the convention established in Fig. 1. See Materials and Methods for details.

**Movie S2. Conformational change between TnsC filaments in the major configuration of transpososome and helical TnsC.** The movie shows the morphing between twelve protomers of TnsC from the major configuration of transpososome and helical TnsC structure (PDB: 7M99).

**Fig. S1.**
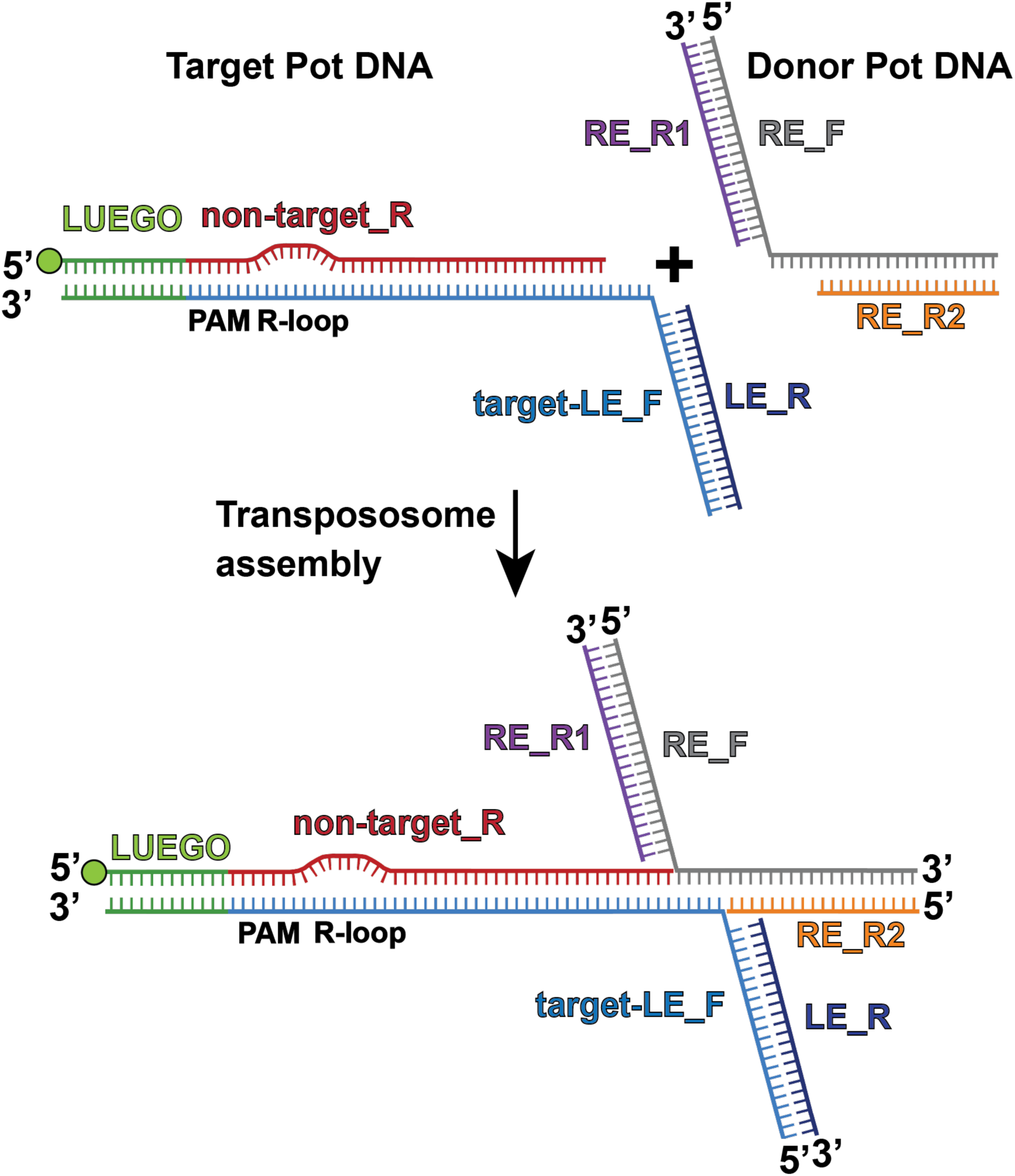
Designed DNA substrate schematic. DNA substrate design for the transpososome assembly includes two DNA substrates: target pot DNA containing a Cas12k-binding site and the first two TnsB binding sites of left-end (LE), and donor pot DNA with the first two TnsB binding sites of right-end (RE). DNA substrates were designed to form a strand-transfer complex by having 5 base pair (bp) single-stranded DNA (ssDNA) connecting the target DNA region and the transposon DNA region. The design for target-pot DNA is composed of four single-stranded DNA (ssDNA) of the following: Target-LE_F (light blue), non-target_R (red), desthiobiotinylated LUEGO (green), and LE_R (dark blue). Cas12k-binding region contains PAM and a 10 base-pair (bp) mismatch to facilitate R-loop formation as indicated as a displaced strand. 5’ end labeled desthiobiotin (green circle) on the LUEGO was used to conjugate target-pot DNA on magnetic beads for the pulldown. Second, donor-pot DNA consists of three ssDNA: RE_F (grey), RE_R1 (purple), and RE_R2 (orange). LE_R region and RE_R1 region of each DNA substrate corresponds to the first two TnsB binding sites of LE and RE, respectively. Two DNA substrates have 5 bp of complementary sequences to each other, which are annealed upon transpososome assembly. Locations of PAM, and R-loop are annotated in black. Sequences for all DNA substrates are included in Table S2.

**Fig. S2.**
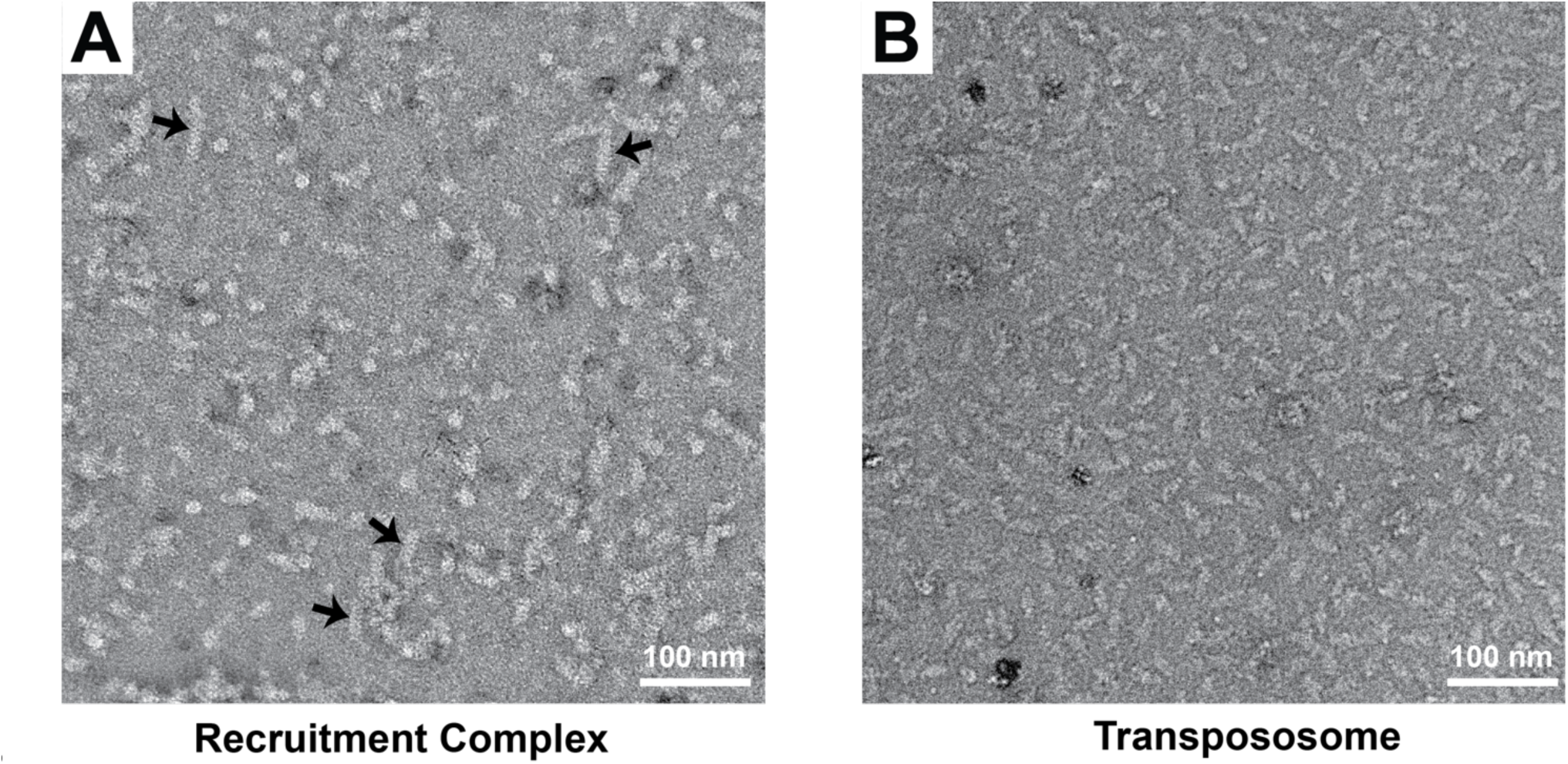
Negative staining electron microscopy shows highly heterogeneous assembly of the target pot, which becomes much more homogeneous upon addition of the donor pot. **A.** Negative stain image of the target pot (i.e. recruitment complex containing Cas12k, S15, TniQ, and TnsC) shows a heterogenous assembly with variable length TnsC filaments (indicated by black arrows). **B.** Negative stain image of the transpososome complex shows the addition of donor pot disassembled TnsC filaments and resulted in a homogeneous sample. Scale bar (white) represents 100 nm.

**Fig. S3.**
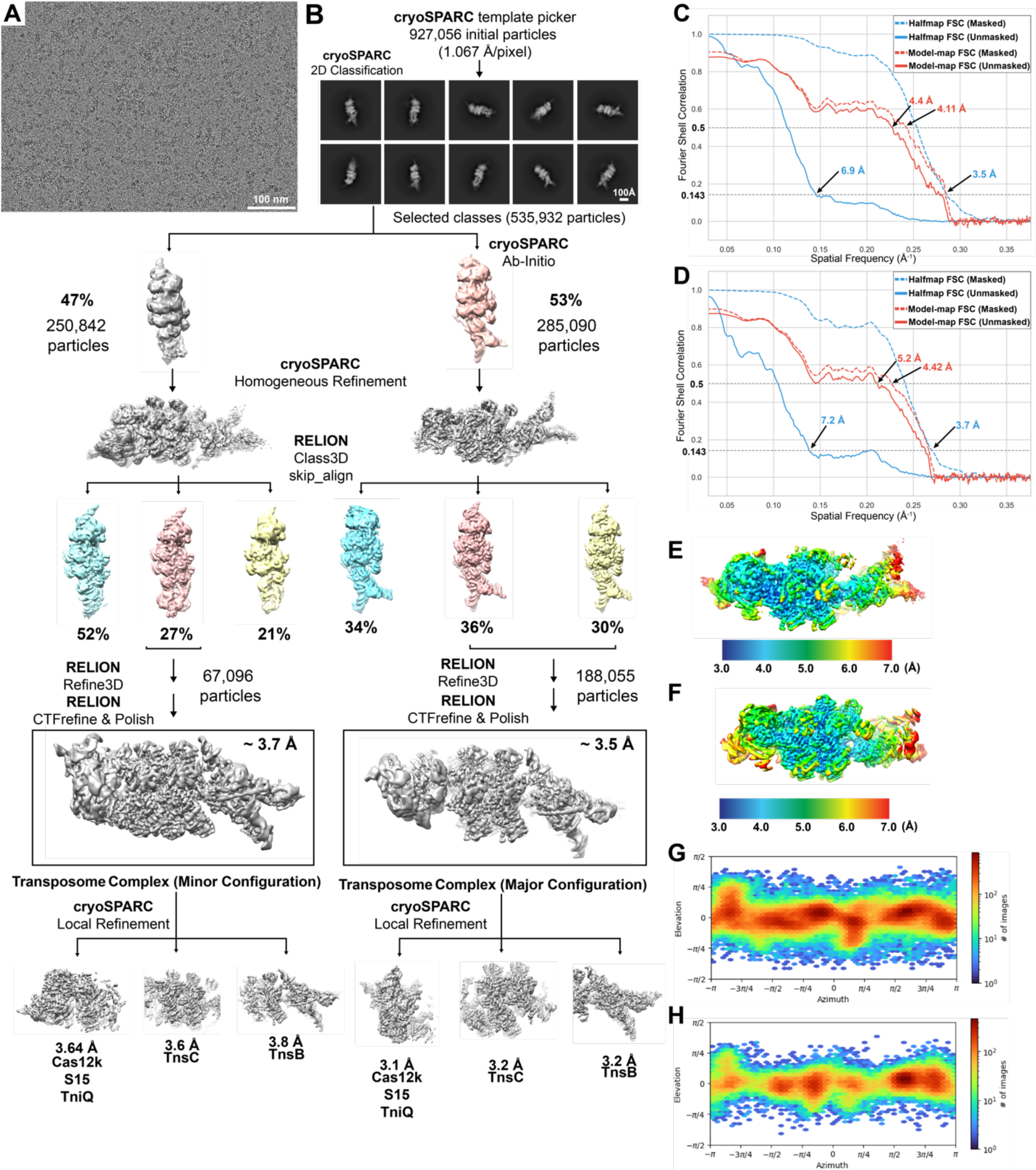
Cryo-EM imaging and image processing pipeline of the transpososome complex. **A.** Representative cryo-EM micrograph from the reconstituted transpososome sample. Scale bar (white, bottom right) represents 100 nm. **B.** Image processing workflow used to analyze the cryo-EM data. 2D classification in cryoSPARC on template picked particles (from 14,017 micrographs) resulted in 535,932 particles. *Ab-initio* reconstruction on the initial particle stack resulted in two classes, one with 53% of the particles (pink) and the other with 47% of the particles (gray). The two classes were separated for subsequent classification and refinement steps. Before performing 3D classification in RELION, each class underwent homogenous refinement in cryoSPARC (*10*). RELION 3D classification (without alignments, skip_align) (*11, 12*) was applied to both populations from the *ab-initio* reconstruction resulting in the colored volumes shown (blue, pink, and yellow). On the left, the two classes (pink and yellow) that have the best resolved Cas12k density were combined for downstream refinement to produce the final 3D reconstruction (boxed), which is the major configuration of the transpososome complex with 12 TnsC protomers. Local refinement was performed on three different segments of the map, focusing on: Cas12k (3.1 Å), TnsC (3.2 Å) and TnsB (3.2 Å). On the right, the class that has the best resolved Cas12k and TnsB density (27% of particles, shown in pink) was selected for downstream refinement to produce the final 3D reconstruction (boxed), which is the minor configuration of the transpososome complex with 13 TnsC protomers. Similar local refinement was performed on three different segments of the minor TnsC complex to result in high resolution reconstructions of the target site proteins (Cas12k+TniQ+S15), TnsC, and TnsB. **C-D.** Fourier shell correlation (FSC) curve of the major (**C**) and minor (**D**) configuration of the transpososome complex, respectively. Masked (dashed) or unmasked (solid) gold standard half-map FSC (blue) and model-map FSC (red) curves are shown for the refined reconstruction and atomic model. Model-map cutoff (0.5) and gold-standard FSC cutoffs (0.143) are indicated with dashed lines. Estimated resolution based on these cutoffs are indicated. **E-F.** Local resolution filtered reconstruction for the major (**E**) and minor (**F**) configuration of the transpososome complex, respectively, are shown with estimated local resolution indicated using colored surface. Local resolution ranges from 3.0 Å (blue) to 7.0 Å (red). Legend at the bottom indicates local resolution range and values in Angstrom. **G-H.** Angular distribution plot for particle projections of the major (**G**) and minor (**H**) configuration of the transpososome complex, respectively. The plot was calculated in cryoSPARC and shows the number of particles for each viewing angle. Colors indicate counts; red corresponds to high particle counts for that particular viewing angle, blue to low particle counts.

**Fig. S4.**
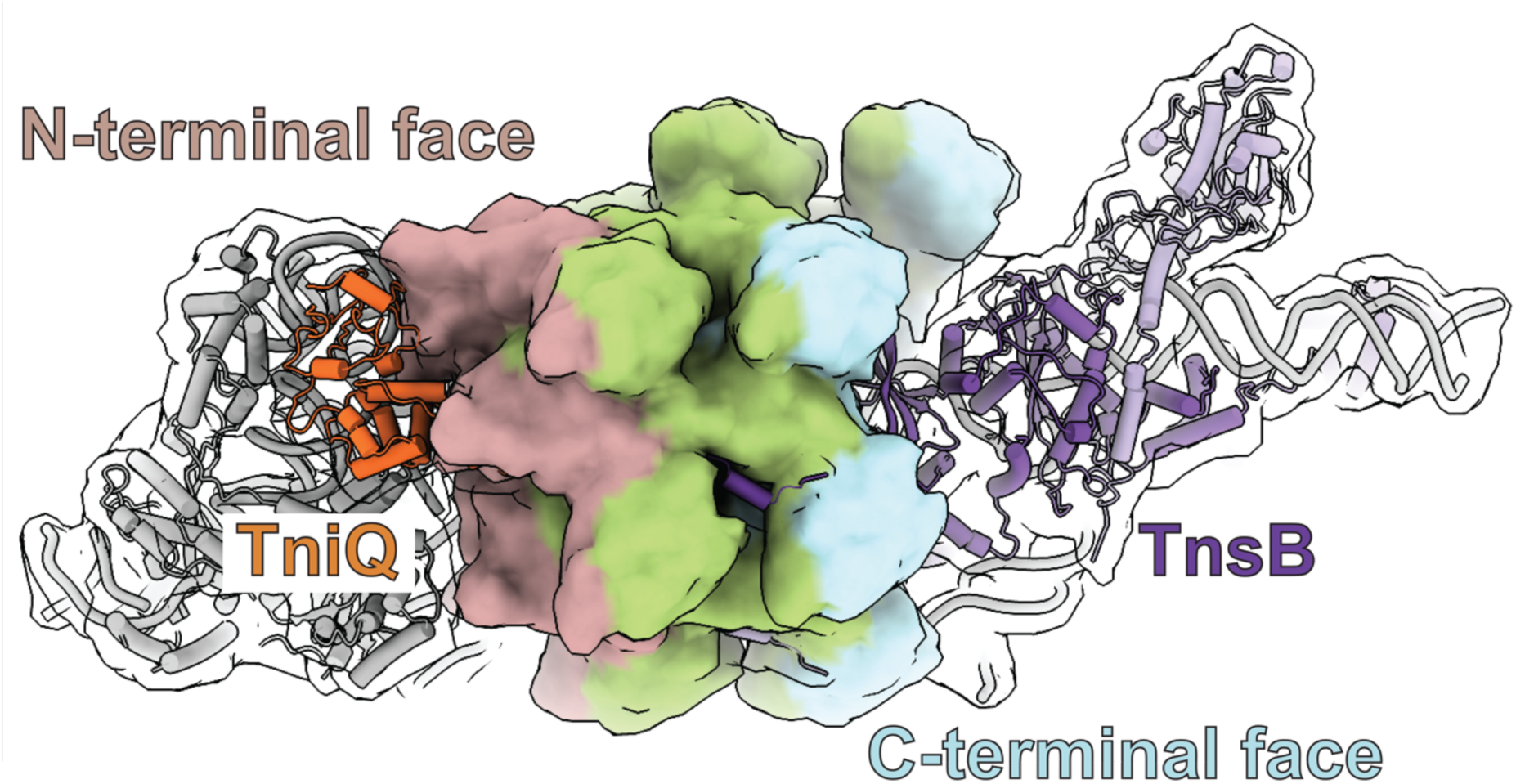
TnsC has dedicated faces for TniQ and TnsB within the transpososome. A simulated map is colored by different regions of TnsC (opaque surface) to represent the N- and C-terminal face of TnsC. The regions are colored as follows: N-terminal face (residues 19-140 on TnsC7-TnsC12, rose-brown), and C-terminal face (residues 141-275 on TnsC1-TnsC6, light blue). Region not included in either N- or C-terminal face is colored green. TniQ (orange ribbon) and TnsB (purple ribbon) associates with the N-terminal face and C-terminal face respectively.

**Fig. S5.**
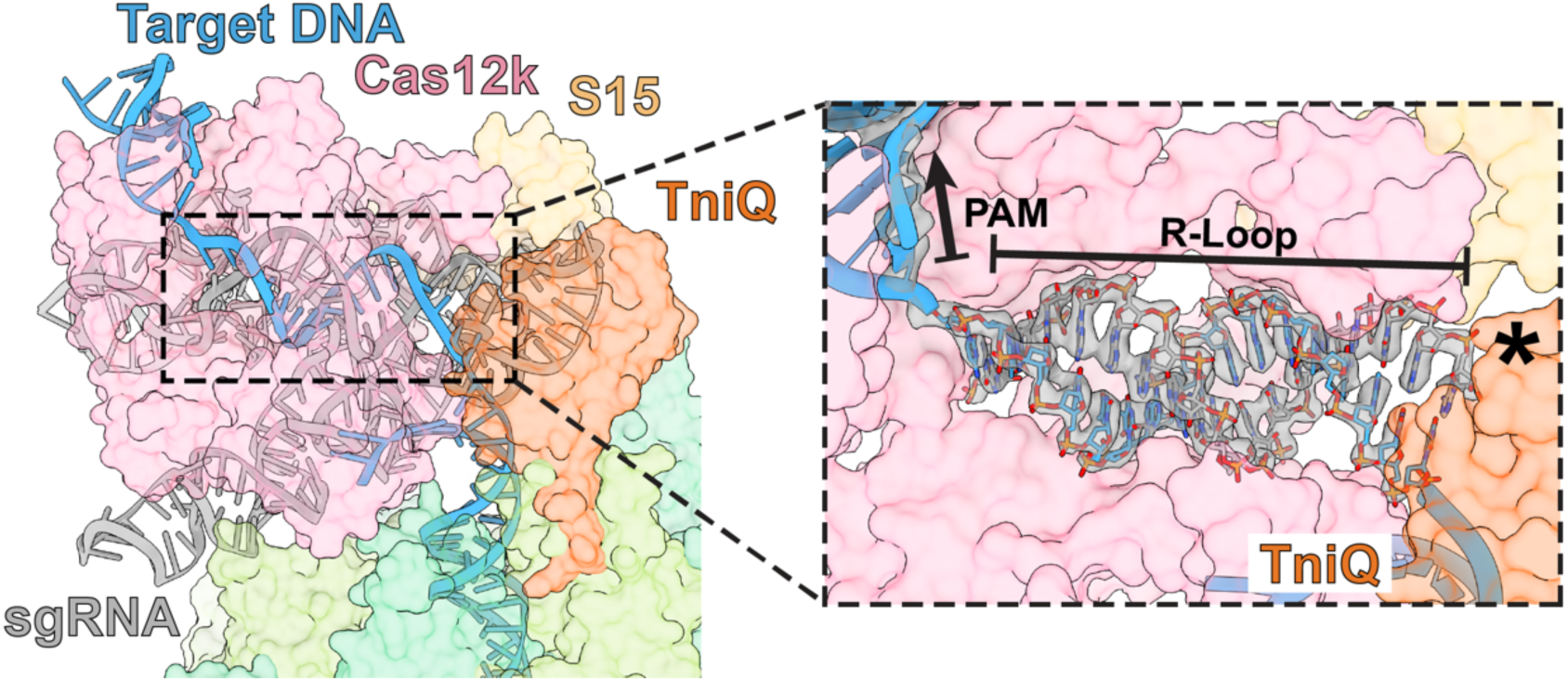
TniQ is located at the PAM-distal end of the Cas12k R-loop. DNA (blue ribbon) forms 17 base pair heteroduplex with RNA (gray) in Cas12k (pink). Cas12k, S15 (tan), and TniQ (orange) are shown in surface representation. Close-up view on the right shows the model docked into the cryo-EM density of the R-loop. The PAM distal end of the R-loop is indicated with an asterisk (*).

**Fig. S6.**
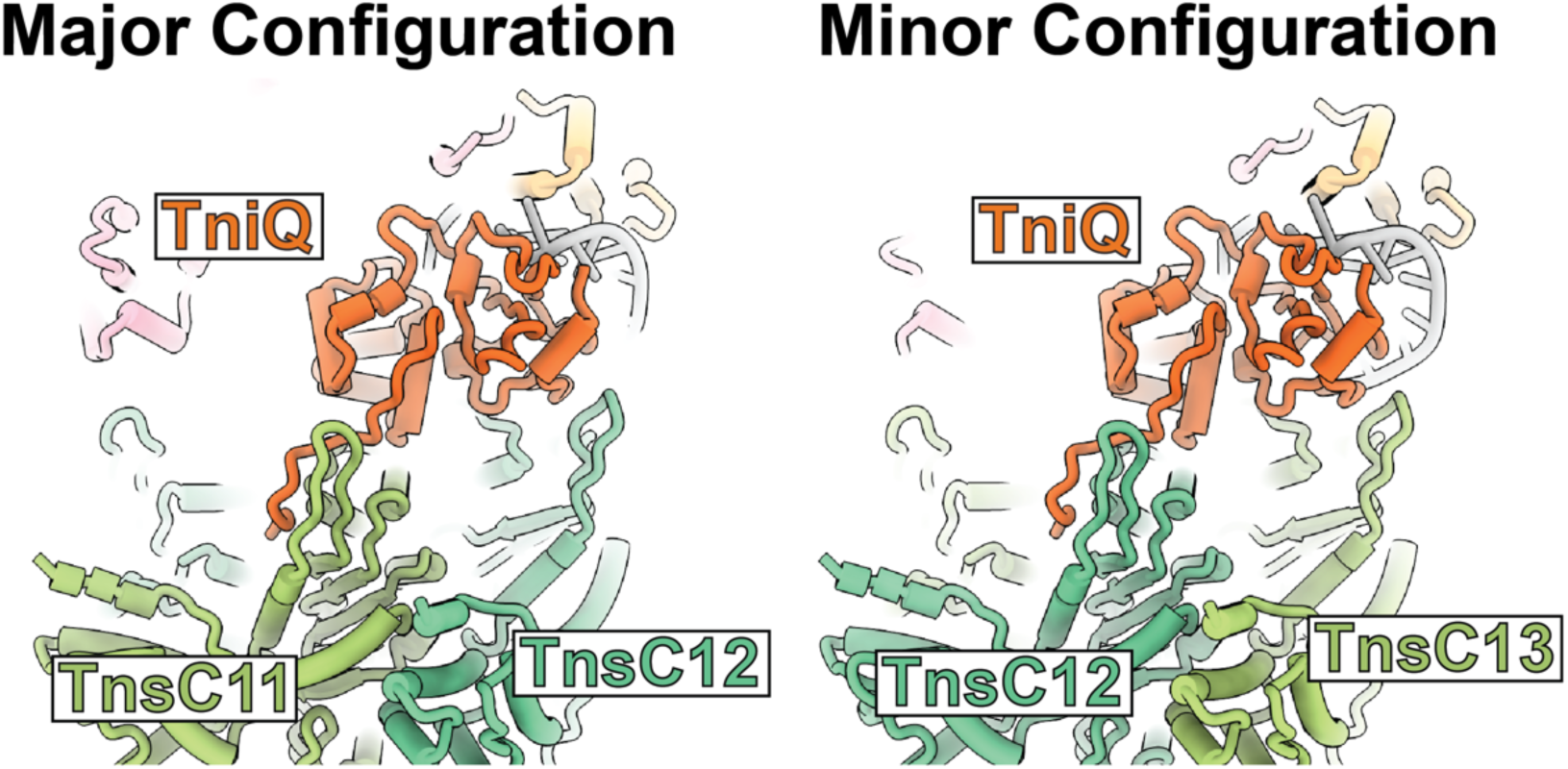
TniQ interacts with two TnsC protomers proximal to Cas12k in both major and minor configurations. In both major (left) and minor (right) configurations, TniQ bridges the two TnsC protomers closest to Cas12k in an identical manner. Colors for the ShCAST components are the same as previously defined in Fig. 1.

**Fig. S7.**
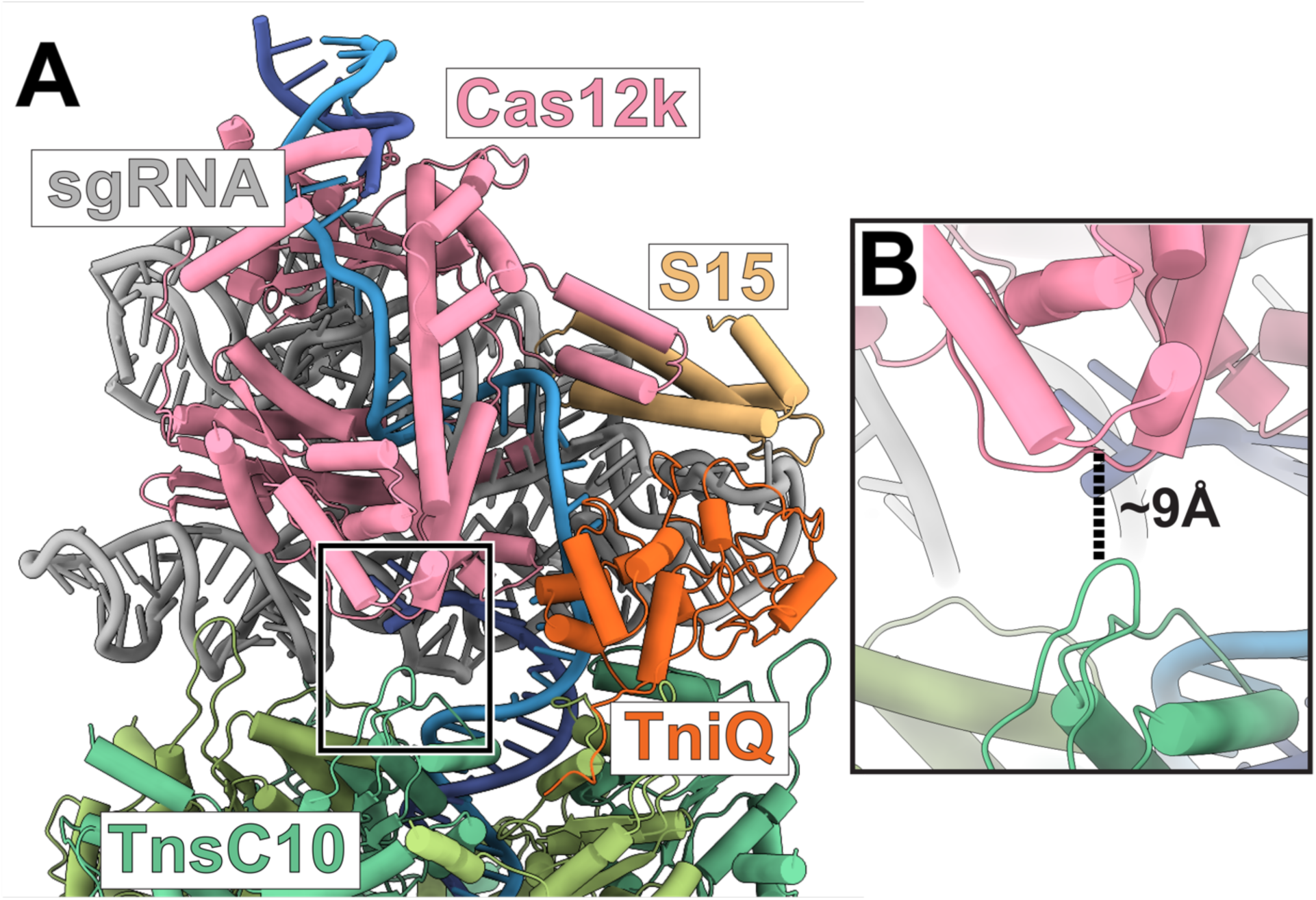
Cas12k is too far away to interact with TnsC. **A.** Atomic model of the ShCAST transpososome, focusing on Cas12k. The TnsC protomer closest to Cas12k (TnsC10) is labeled. TnsC protomers are numbered as previously defined (see Main Text, Fig. 2). ShCAST protein and nucleic acid components are labeled and colored according to previously defined colors (see Main Text, Fig. 1). Black box indicates the TnsC finger loop that is close to Cas12k shown as inset in panel B. **B.** The closet distance between TnsC protomer TnsC10 and Cas12k is shown with dashed line and labeled (in Å).

**Fig. S8.**
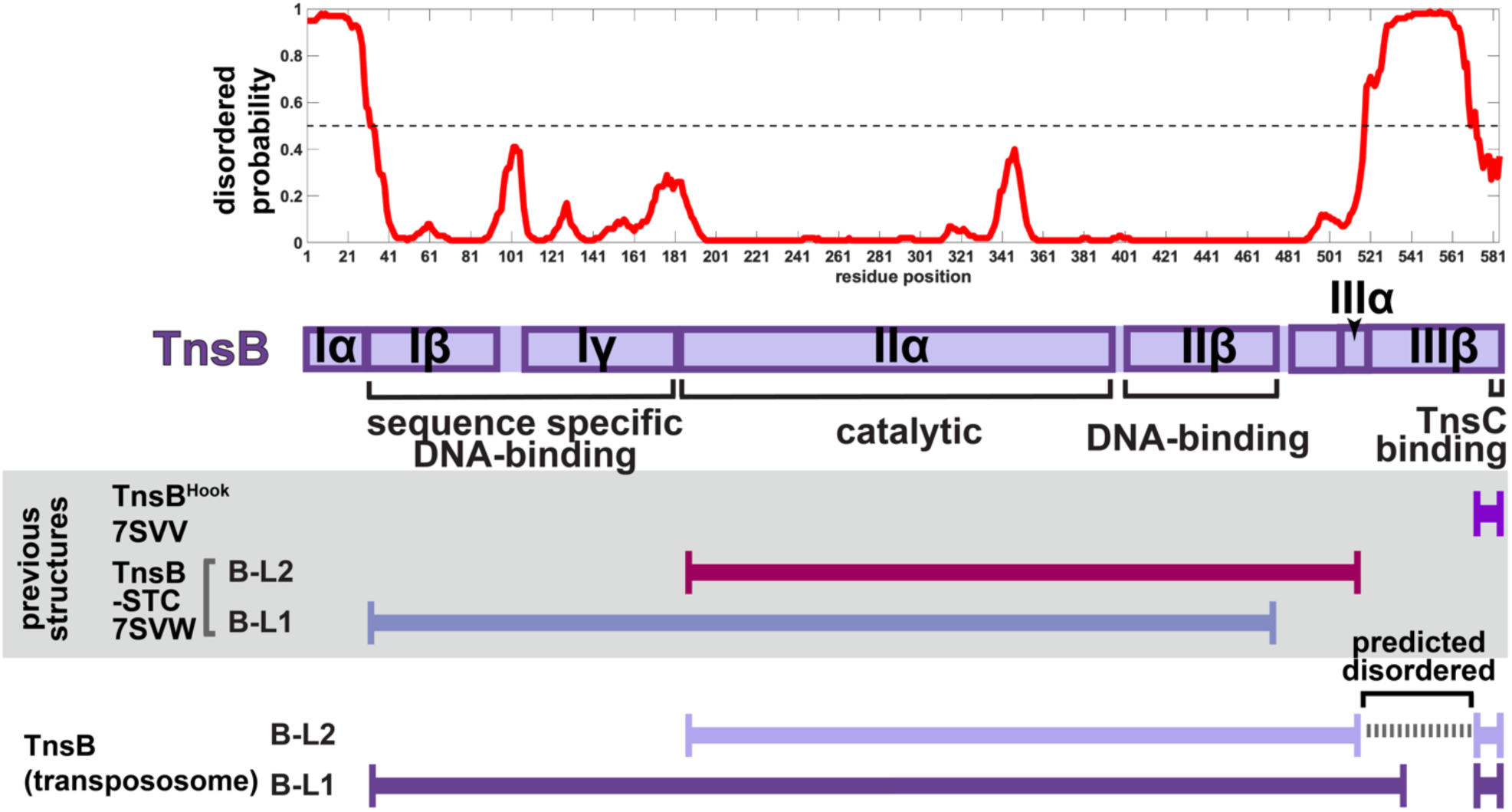
TnsB disordered prediction and modeled segments. Top panel: The probability of disorder at each residue position is plotted. 1 indicates high probability of disorder, 0 indicates low probability of disorder. Dashed line indicates 50% probability, the cutoff above which to consider a residue position to be predicted disordered. The purple diagram below the disordered prediction is the domain diagram of TnsB, along with the assigned functions for each domain. Domain names follow that of MuA. Gray box indicates the previous structures of TnsB resolved; The TnsB^Hook^ peptide corresponds to the last fifteen residues of TnsB, 570-584 (PDB: 7SVV). B-L1 and B-L2 are the two named conformations of TnsB resolved in the strand-transfer complex reported previously (PDB: 7SVW), also solved by cryo-EM. Bottom panel indicates the regions of B-L1 and B-L2 resolved within the transpososome. The stretch of residues corresponding to the flexible linker connecting the structured core of TnsB to the C-terminal TnsB^Hook^ is indicated in dashed gray lines.

**Fig. S9.**
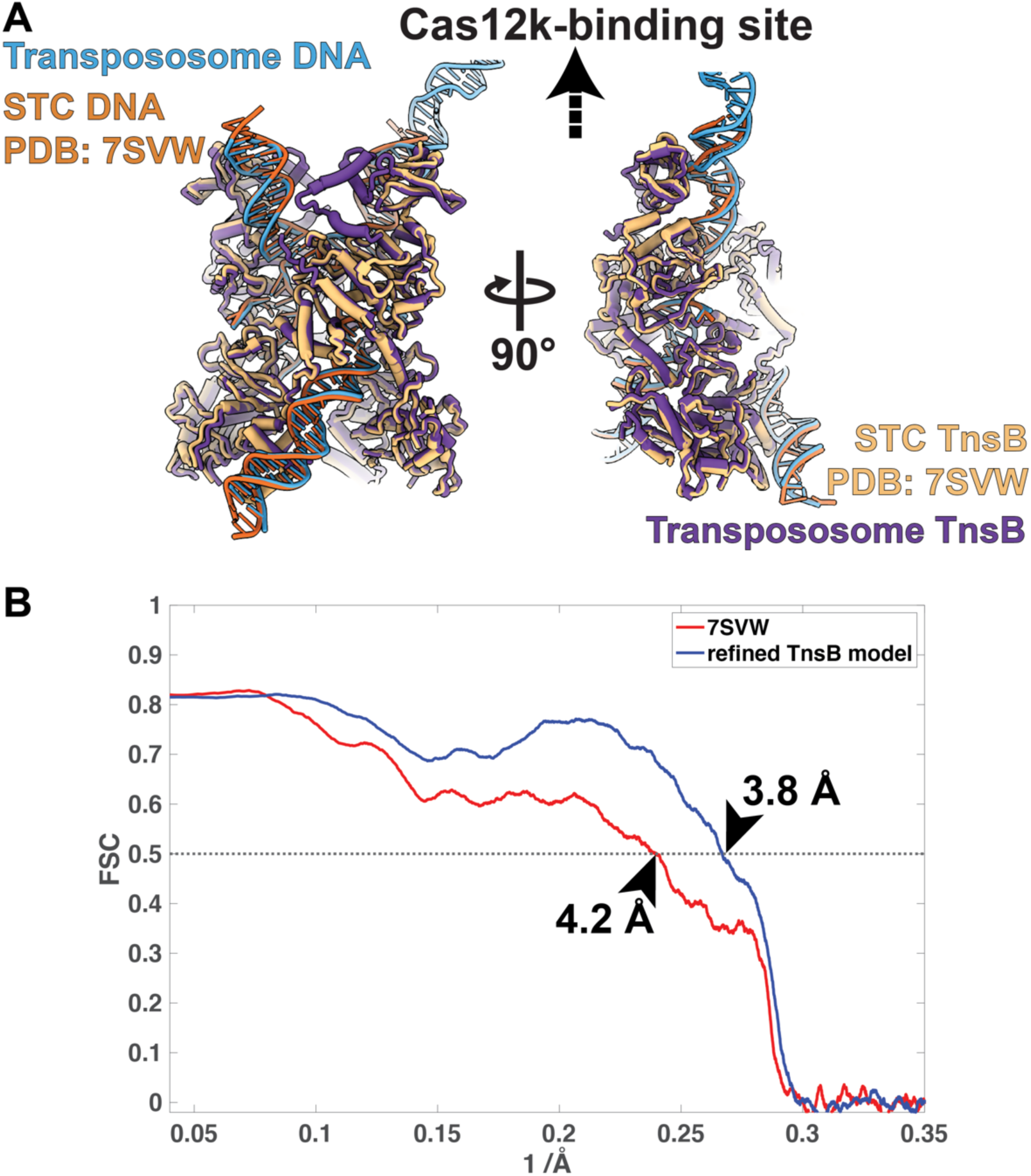
TnsB structure in the ShCAST transpososome is approximated well by the previously determined TnsB strand-transfer complex. **A.** The refined TnsB atomic model (this paper, purple) is superimposed onto the previous TnsB strand-transfer complex (STC) shown in tan (PDB: 7SVW). Overall architecture of the complexes is identical with the Cα r.m.s.d. of 1.54 Å. Target-DNA and transposon DNA (light blue) also appear structurally similar to the TnsB STC (orange). Two different orthogonal views are shown for comparison. The direction of the Cas12k-bound target-site is indicated with arrow. **B.** The masked model-map Fourier-shell correlation (FSC) curves are shown for the rigid-body docked TnsB STC (PDB: 7SVW, red curve) and the refined TnsB model (blue curve). A cutoff of 0.5 (dashed line) indicates the estimated resolution, or ‘goodness of fit’ of the atomic model against the map. The map used here is the locally refined TnsB cryo-EM reconstruction. Estimated resolution of each model is indicated with arrows. As expected, molecular refinement improved upon the starting model (PDB: 7SVW) from 4.2 Å to 3.8 Å, and roughly corresponds to the resolution of the locally refined cryo-EM map (3.2 Å).

**Fig. S10.**
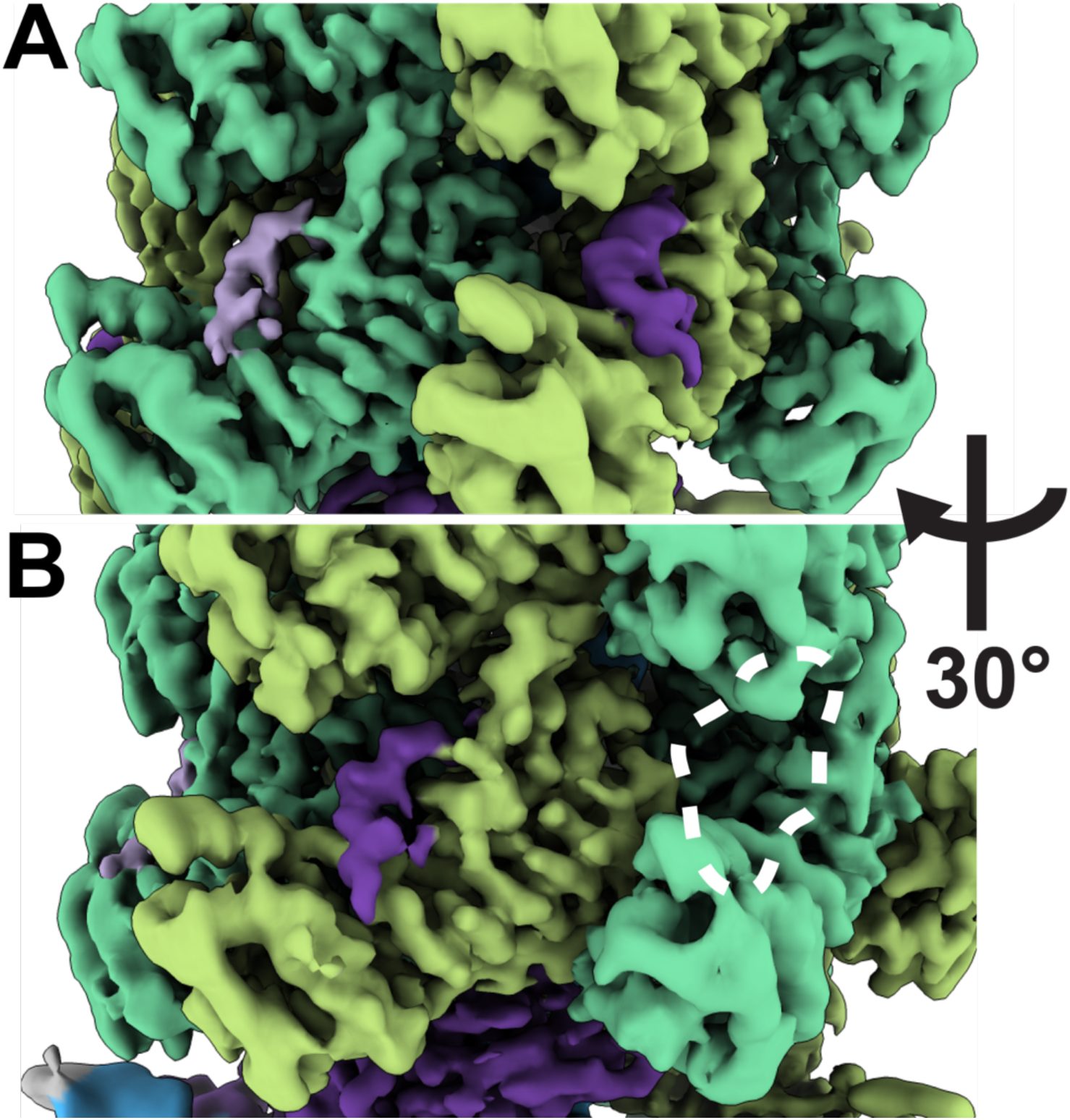
TnsB^Hook^ density occupies select TnsC protomers in the TnsC hexamer closest to TnsB. **A.** Local resolution filtered cryo-EM reconstruction (same as that shown in Fig. 1) colored by assignment reveals that TnsB^Hook^ (light and dark purple) occupies binding sites on TnsC (green). **B.** Rotation of the cryo-EM reconstruction by 30° shows the adjacent TnsB^Hook^ binding pocket on TnsC (white dashed lines) is empty.

**Fig. S11.**
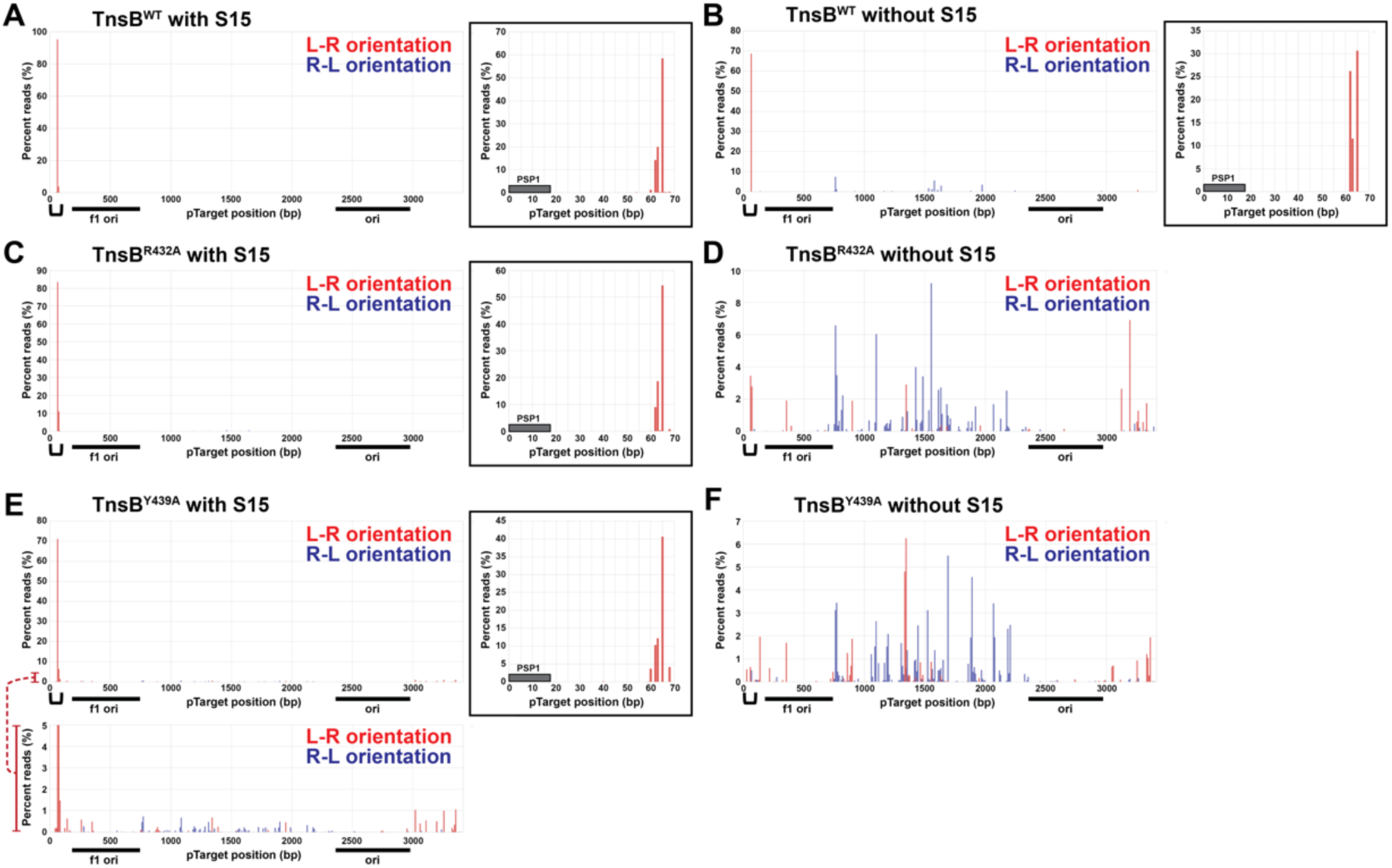
High throughput mapping of the *in vitro* transposition events reveals that the identified interaction between TnsC and TnsB IIΔ domain is crucial for target-site selection. Insertion positions were determined by Illumina sequencing of the plasmids extracted from colonies from *in vitro* transpositions under the following conditions (same with Fig. 3F, see Materials and Methods): **A.** TnsB wild-type (WT) with S15, **B.** TnsB WT without S15, **C.** TnsB R432A with S15, **D.** TnsB R432A without S15, **E.** TnsB Y439A with S15, and **F.** TnsB Y439A without S15. Determined insertion positions were plotted as a histogram indicating the percentage of the reads at the 10 base-pair (bp) windows within the target plasmid. Position numbers (x-axis) correspond to the number of base pairs between the PAM and the beginning of the transposon-end sequence after the transposition. Red and blue bars represent the transposition products with the left end-right end (L-R, correct) or the right end-left end (R-L, wrong) orientation respectively. For the conditions with high on-target percentage (> 60%, panels A, B, C, and E), insets are presented for the positions around the PSP1 protospacer (from 0 bp to 70 bp), which is indicated with black brackets on the x-axis. Grey bar in the inset indicates the 17 bp PSP1 protospacer. For panel E, a red bar on the y-axis represents the region for zoom-in on the lower panel to visualize signals from the off-target transposition events. Two origins of replications within the target plasmids (f1 ori and ori) are annotated as black bars on the x-axis, which explains the reason for the cold spots for the transpositions.

**Fig. S12.**
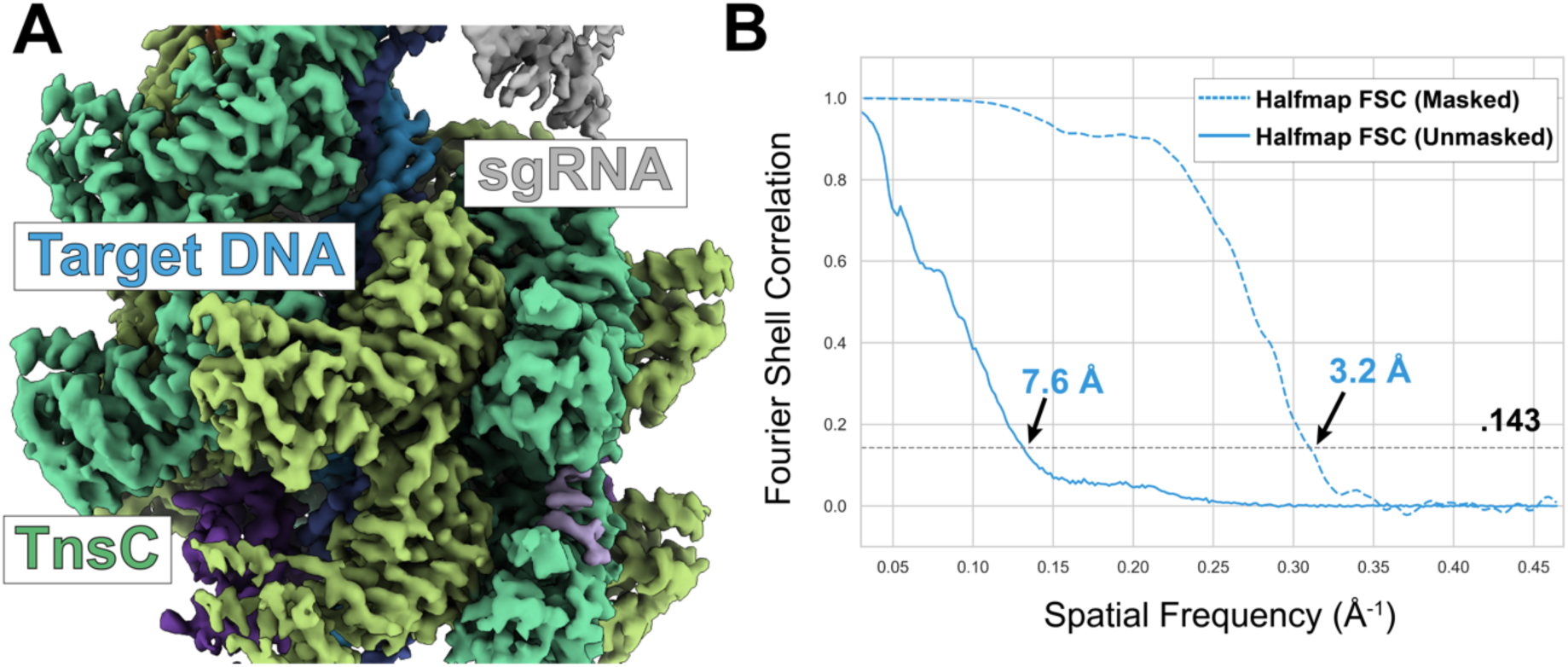
High-resolution cryo-EM reconstruction of TnsC local refinement. **A.** The local refinement cryo-EM map focused on the TnsC region of the ShCAST transpososome, colored according to previously defined colors (see Main Text, Fig. 1). The sections of the transpososome that are not included in the refinement mask, including sgRNA (grey), are by comparison less well resolved. **B.** Fourier shell correlation (FSC) curve of the local refinement cryo-EM reconstruction of the TnsC region of the major configuration. Masked (dashed) or unmasked (solid) gold standard half-map FSC (blue) are shown for the refined reconstruction. Gold-standard FSC cutoff (0.143) is indicated with dashed line and estimated resolution based on this cutoff is indicated.

**Fig. S13.**
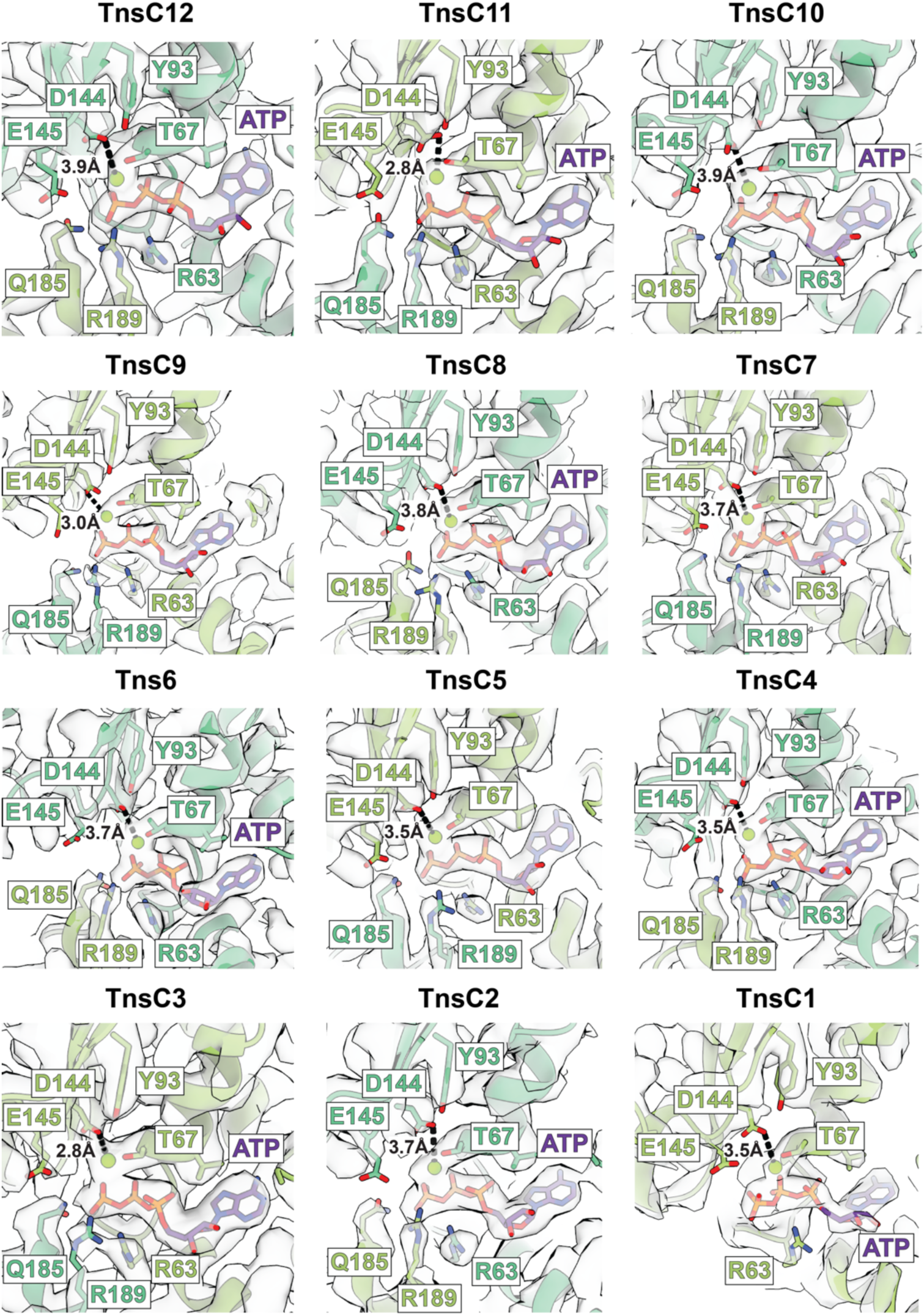
ATP is bound at all TnsC protomers in the ShCAST transpososome. ATP (purple) is bound at each of the TnsC protomers (numbered 1 through 12) in the ShCAST transpososome. Each of the ATP binding pockets is shown, docked into the cryo-EM density map (transparent gray surface). Magnesium is shown as a green sphere, and relevant sidechains are shown in stick. Because the ATP binding pocket sits between two TnsC protomers, either light green or dark green is used to shade the particular sidechains displayed, depending on which protomer (and therefore which ATP-binding pocket) is being shown. E145 and D144 are the catalytic residues, and T67 coordinates ATP-binding. Q185 and R189 are the sensor residues from the neighboring TnsC protomer that complete the ATP binding pocket. Distance from D144 to Magnesium is labeled in Å, and the interaction is shown with dashed lines.

**Fig. S14.**
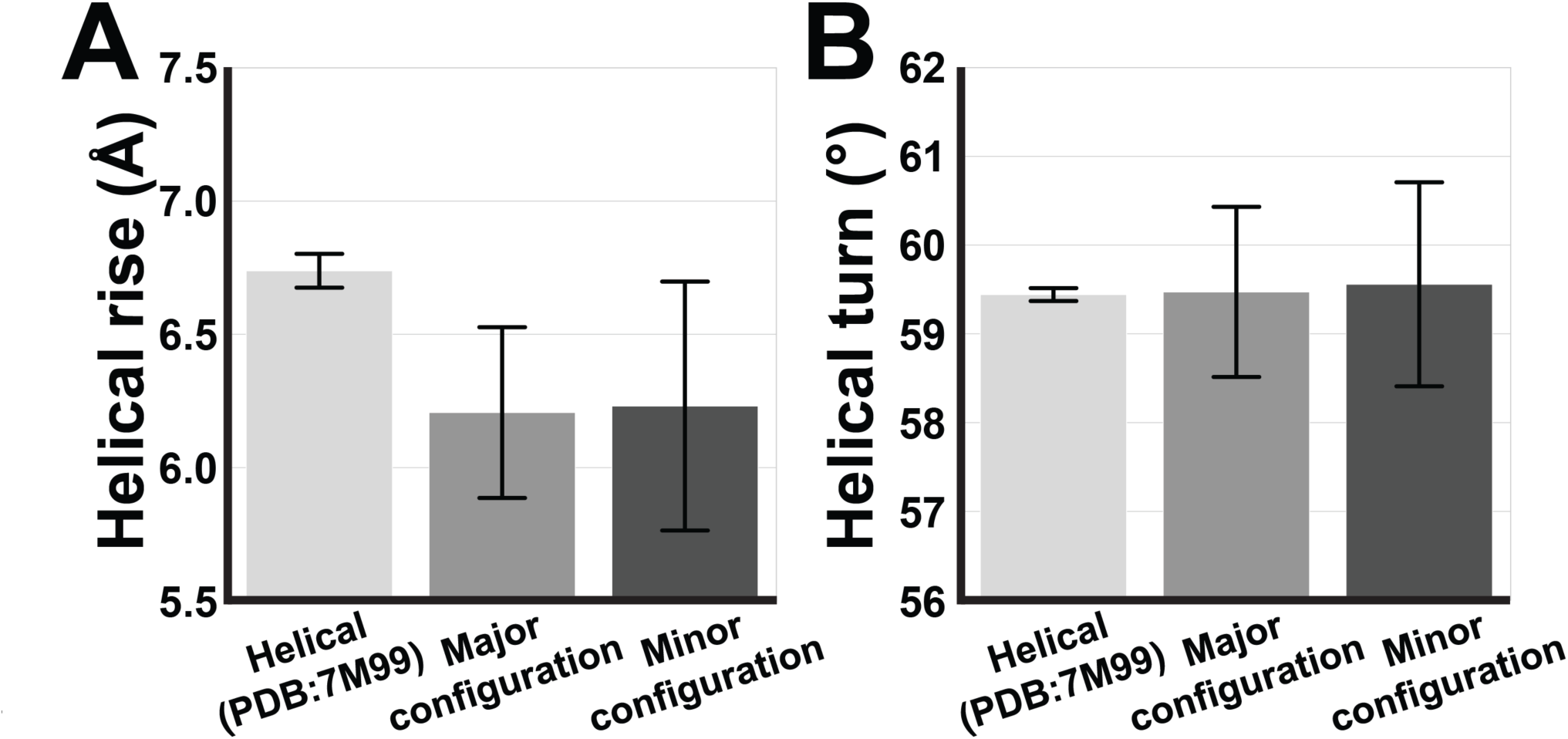
Measured helical parameters of the major and minor configurations of transpososomes are more variable compared to the helical TnsC structure. Average helical rise **(A)** and helical turn **(B)** estimated between TnsC protomers within ATPγS-bound helical TnsC (PDB: 7M99), TnsC from major configuration and minor configuration of transpososome. Helical parameters were estimated using Rosetta suite (see Materials and Methods for detail), and averaged to plot a bar-graph. Error bar indicates standard deviations from the estimated values. Both configurations of the transpososome have lower values of helical rise, and comparable values of helical turn. Large standard deviations of both helical rise and turn suggest that TnsC in context of the transpososome does not follow defined helical symmetry.

**Fig. S15.**
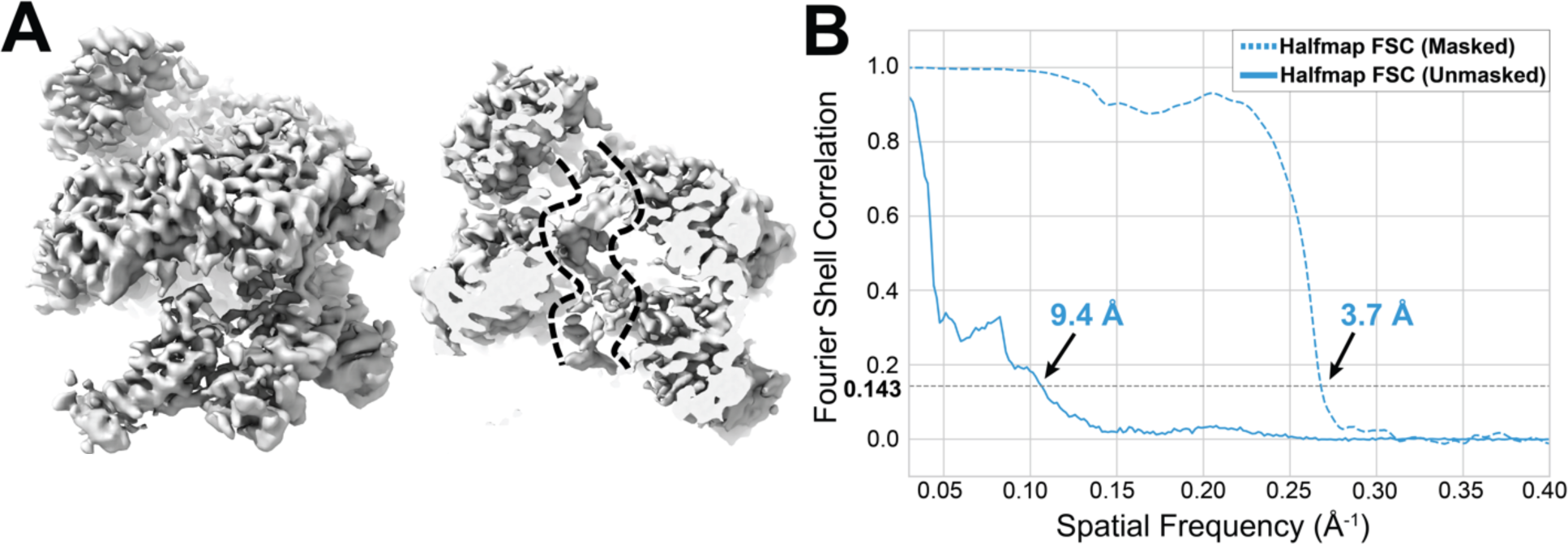
Reconstruction of the two turns of TnsC in transpososome using helical reconstruction approaches results in a lower-resolution map. **A.** Helical Cryo-EM reconstruction of TnsC in the major configuration. Helical parameters of the ATPγS-bound TnsC filaments (rise = 6.82 Å and twist = 60°) were used to impose helical symmetry. Local search of the helical parameters was allowed (see Materials and Methods for detail). The reconstruction (left) yields a lower resolution and a map with aberrant features, where the density of DNA in the map (right, indicated by dashed lines) is barely visible, compared to the result of local refinement (3.2 Å. Fig. S12). These results indicate that the TnsC protomers in transpososome does not follow the helical symmetry of TnsC filaments. **B.** Fourier shell correlation (FSC) curve of the helical refinement cryo-EM reconstruction of the TnsC region of the major configuration. Masked (dashed) or unmasked (solid) gold standard half-map FSC (blue) are shown for the refined reconstruction. Gold-standard FSC cutoff (0.143) is indicated with dashed line and estimated resolution based on this cutoff is indicated.

**Fig. S16.**
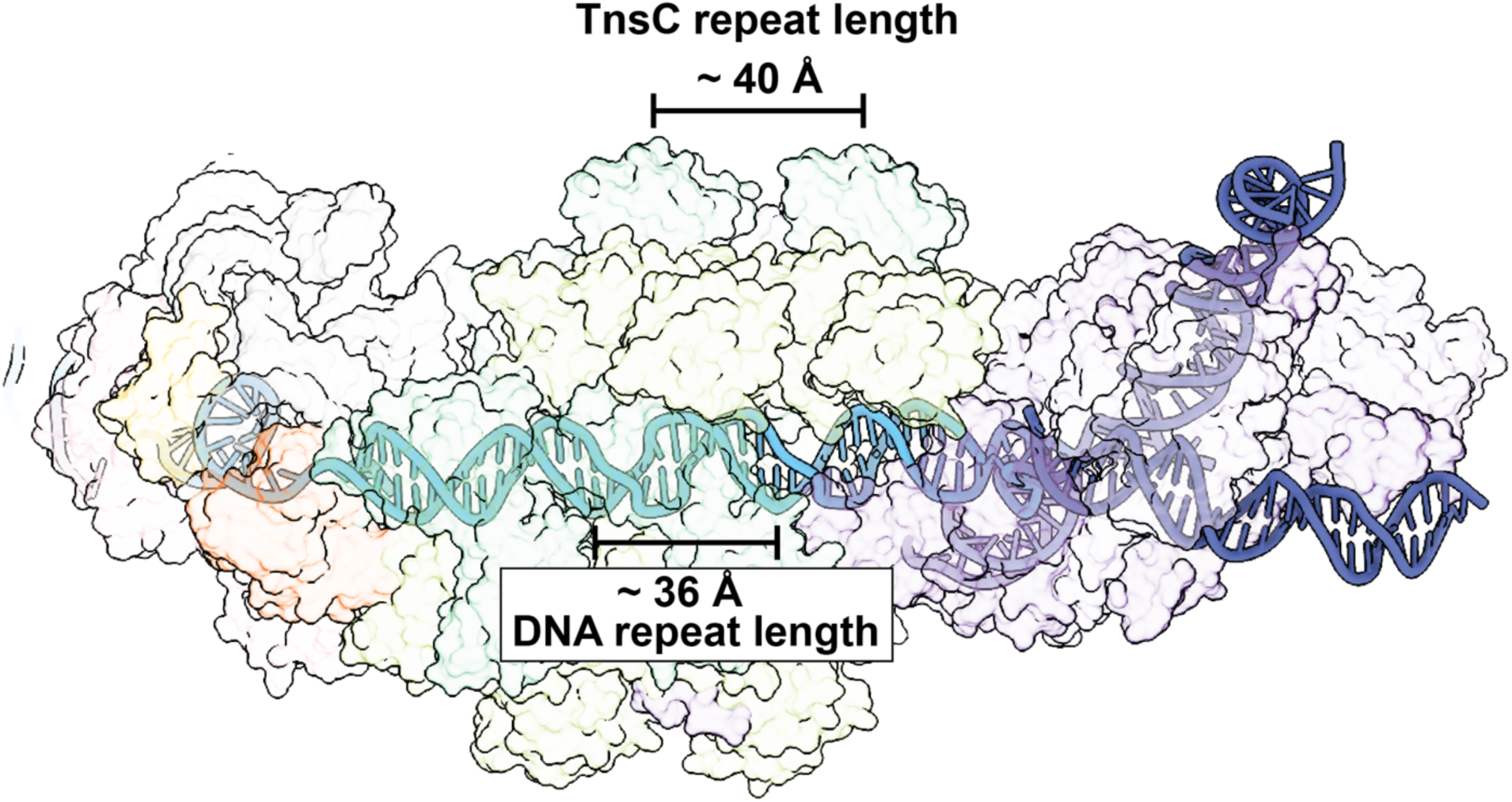
TnsC in ShCAST transpososome does not match helical parameters of the bound DNA. The repeat length of TnsC turn (∼6 protomers per turn) is approximately 40 Å, while the repeat length of the TnsC-bound DNA (∼11 base pairs per turn) is approximately 36 Å. DNA model is represented in a solid ribbon. Protein components and sgRNA in the transpososome are represented as transparent surfaces. Color scheme is identical to the established colors in Fig. 1.

**Fig. S17.**
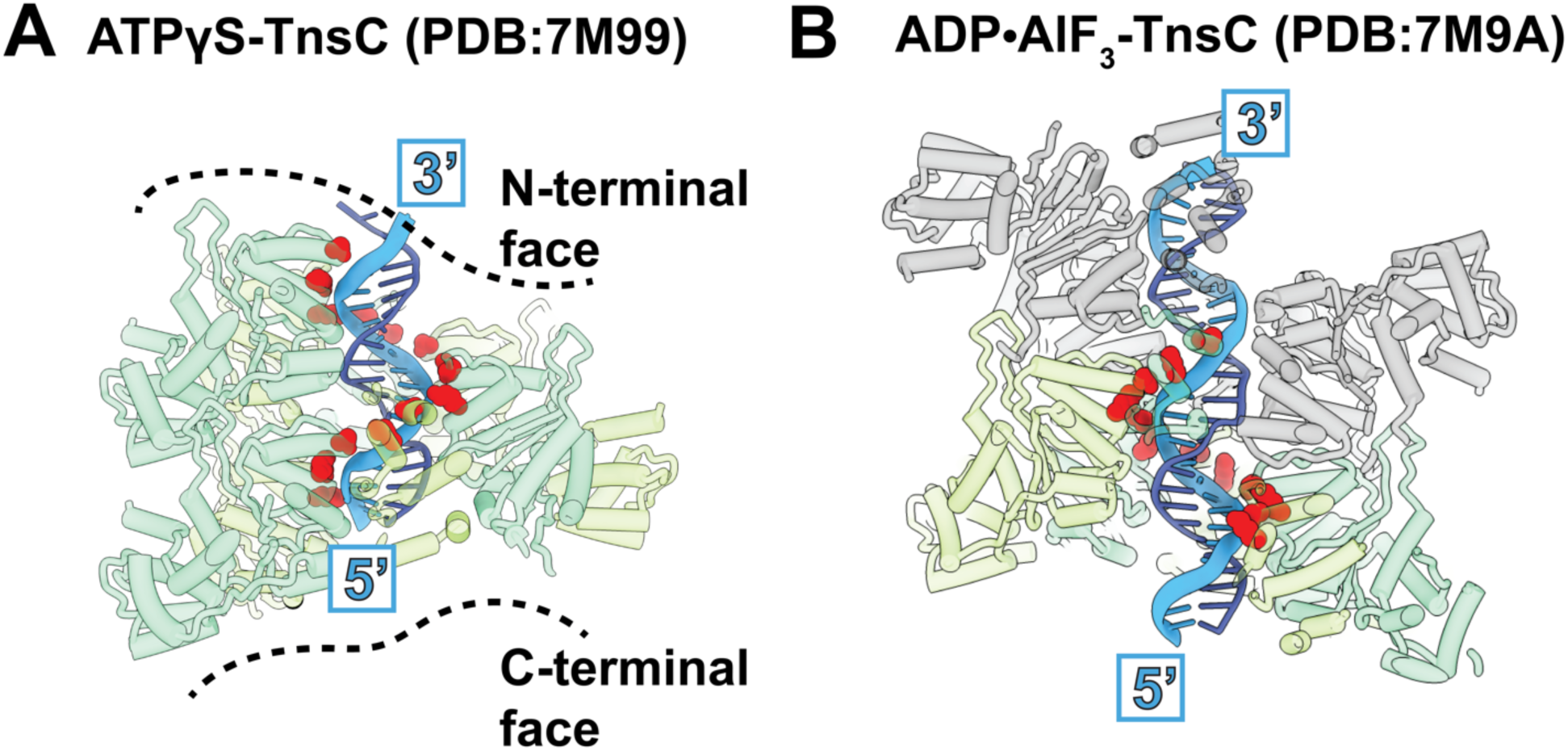
Previous TnsC helical filament structures selectively interact with one strand of DNA. **A.** The previously determined structure of ATPγS bound-TnsC (PDB: 7M99) is shown. TnsC (green) is bound to the DNA duplex (blue). Importantly, TnsC residues K103 and T121 (red spheres) track one strand of DNA (light blue, shown in thicker width) in the 5’ to 3’ direction going in the direction of TnsC’s C-terminal to N-terminal face (dashed black lines). The 5’ and 3’ end of the DNA strand interacting with TnsC are labeled. **B.** ADP·AlF_3_ bound-TnsC (PDB: 7M9A) shows two hexamers, oriented in opposite directions. For clarity, the hexamer shown in the same orientation as in panel A is shown in green, and the other hexamer is shown in gray. DNA strand interacting with the green TnsC protomers is represented in light blue with thicker width.

**Fig. S18.**
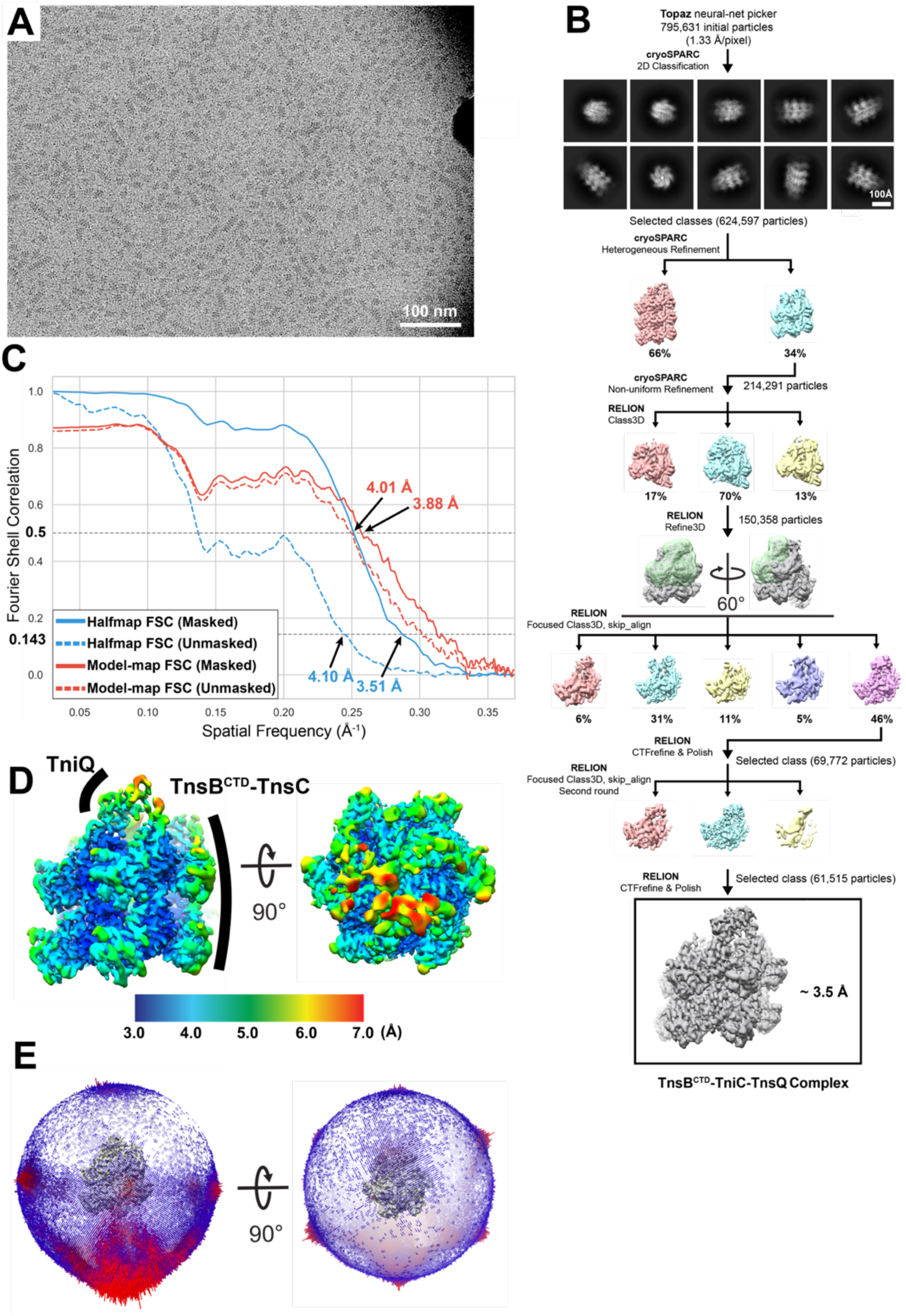
Image processing pipeline for TnsB^CTD^-TnsC-TniQ structure. **A.** Representative cryo-EM micrograph of the TnsB^CTD^-TnsC-TniQ sample. TnsB^CTD^ corresponds to TnsB residues from 476 to 584. White scale bar represents 100 nm. **B.** Image processing workflow of TnsB^CTD^-TnsC-TniQ dataset. From initial 2D classification, classes with high-resolution 2D averages were selected for heterogeneous refinement in cryoSPARC(*10*). Class with two turns of TnsC (cyan, 214,291 particles) was subjected to RELION(*11, 12*) 3D classification, which resulted in a class with better cryo-EM density of TniQ (cyan, 150,358 particles). This particle stack was further subjected to two subsequent rounds of focused classification without alignment using the mask (green, overlaid on the grey input reconstruction) that includes one TniQ and two TnsC protomers. This resulted in the final particle stack of 61,515 particles. Final 3D refinement in RELION converged to a final ∼ 3.5 Å reconstruction of the TnsB^CTD^-TnsC-TniQ complex. **C.** Fourier-shell correlation (FSC) curve from gold-standard FSC (blue), and model-map FSC (red). Masked and unmasked FSC were represented in solid and dashed line respectively. The reported resolution was estimated based on the appropriate cutoff (0.143 for half-maps, and 0.5 for model-map FSC), and indicated on the plot. **D.** Local resolution maps from the final reconstruction of TnsB^CTD^-TnsC-TniQ complex. Legend indicates corresponding resolutions of the surface color. TniQ region and TnsB^CTD^-TnsC region of the reconstruction are indicated. **E.** Euler angle distribution of the final reconstruction. Red bars indicate the population of particles in a specific orientation.

**Fig. S19.**
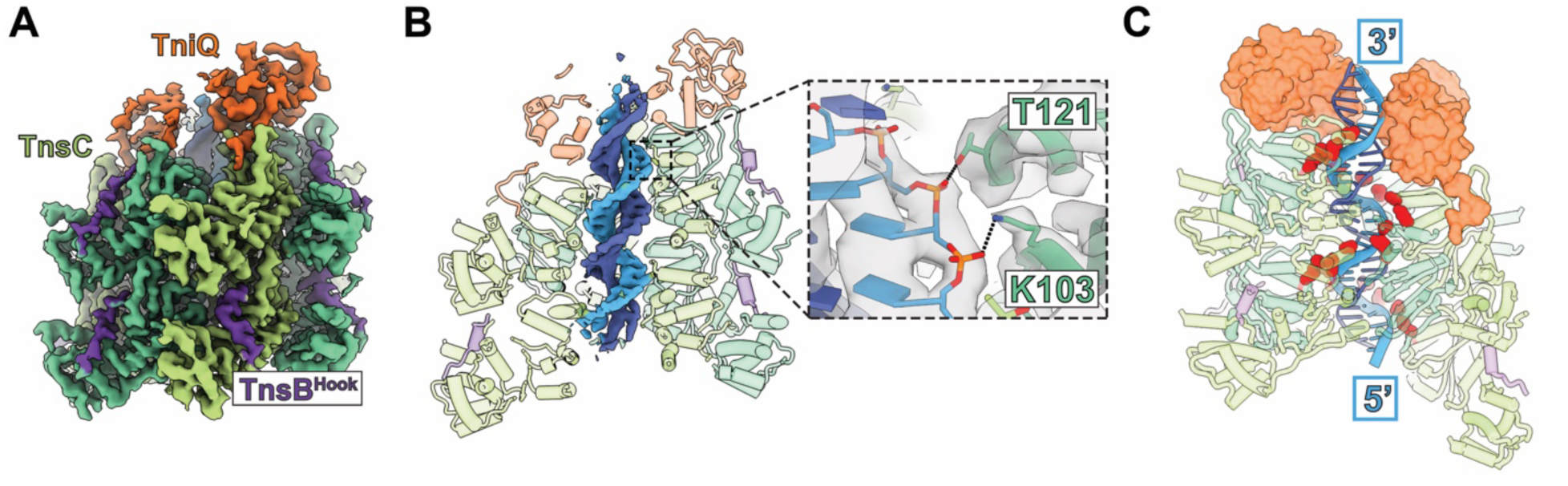
High-resolution cryo-EM reconstruction of TnsB^CTD^-TnsC-TniQ shows TnsC interacts with DNA in the same fashion as previous TnsC structures. **A.** High-resolution cryo-EM structure of TnsB^CTD^-TnsC-TniQ complex. TnsB^Hook^, TnsC, and TniQ are represented in purple, green, and orange, respectively. **B.** Cryo-EM structure of the TnsB^CTD^-TnsC-TniQ complex has high enough resolution to reveal atomic interactions between TnsC (green) and DNA (blue). DNA is shown as segmented density from the full map while TnsB^Hook^, TnsC and TniQ are shown in ribbons. Close-up view on the right shows residues K103 and T121 from TnsC are close enough to form interactions (shown with dashed black lines) with the DNA sugar-phosphate backbone (blue, sticks). Both DNA and the residues fit well into the cryo-EM density (gray, transparent). **C.** TniQ (orange surface) subunits are bound to the N-terminal face of TnsC (green ribbon). Residues K103 and T121 from TnsC interact with one strand of DNA (light blue, shown in thicker width), consistent with previous TnsC structures. Residues K103 and T121 are shown in red spheres. The 5’ and 3’ ends of the DNA strand interacting with TnsC are labeled.

**Fig. S20.**
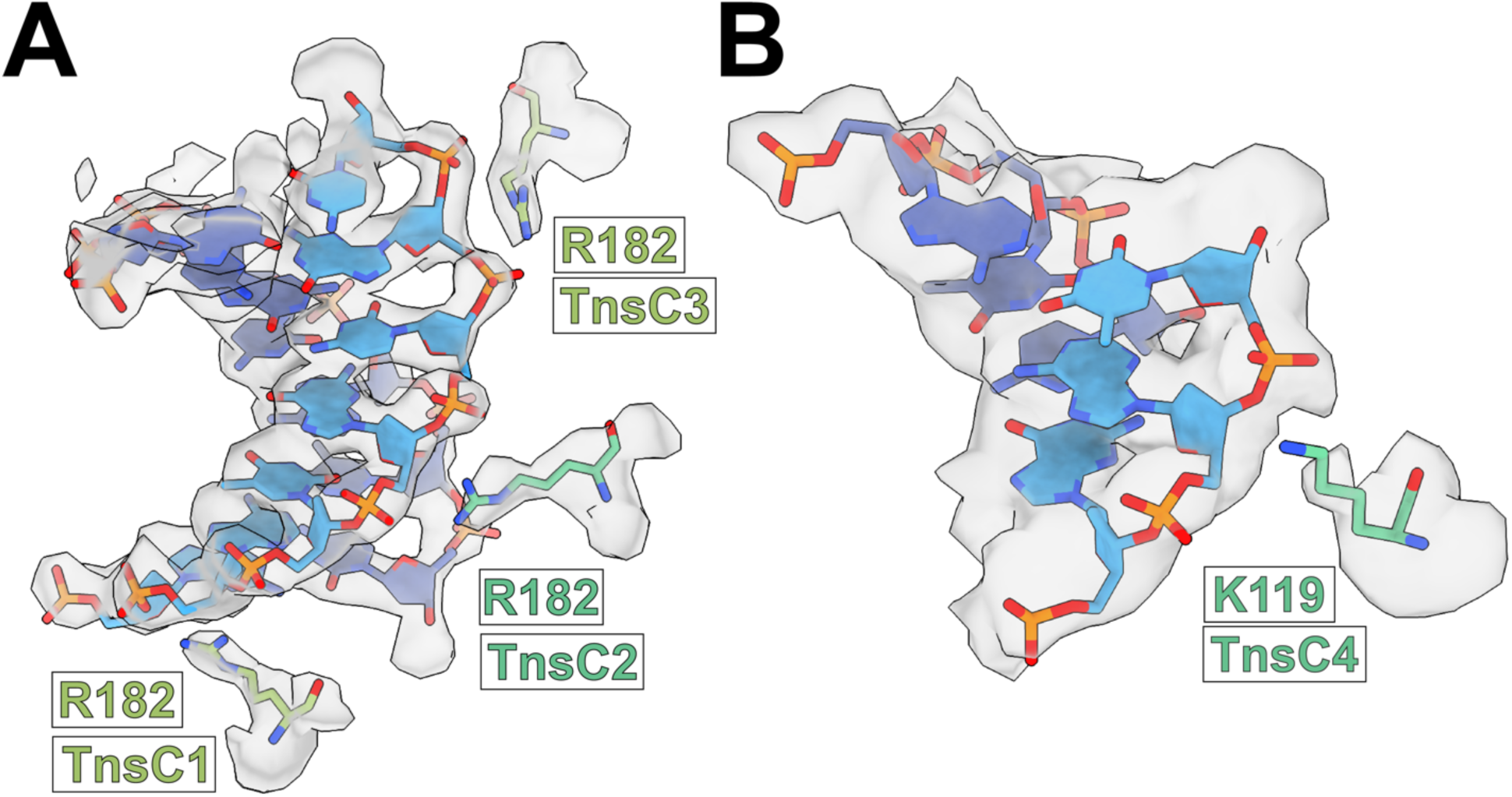
Cryo-EM density supports the newly identified interactions between TnsC and DNA in transpososome. **A.** Residue R182 in TnsC1-TnsC3 is positioned toward the sugar-phosphate backbone of DNA and is supported by cryo-EM density. Zoned map of the TnsC local refinement map is transparent. The map is zoned around the region containing both DNA strands and R182, shown in filled and stick representation, respectively. **B.** Residue K119 in TnsC4 is positioned toward the sugar-phosphate backbone of DNA and is supported by cryo-EM density. Zoned map of the TnsC local refinement map is transparent. The map is zoned around the region containing both DNA strands and K119, shown in filled and stick representation, respectively.

**Fig. S21.**
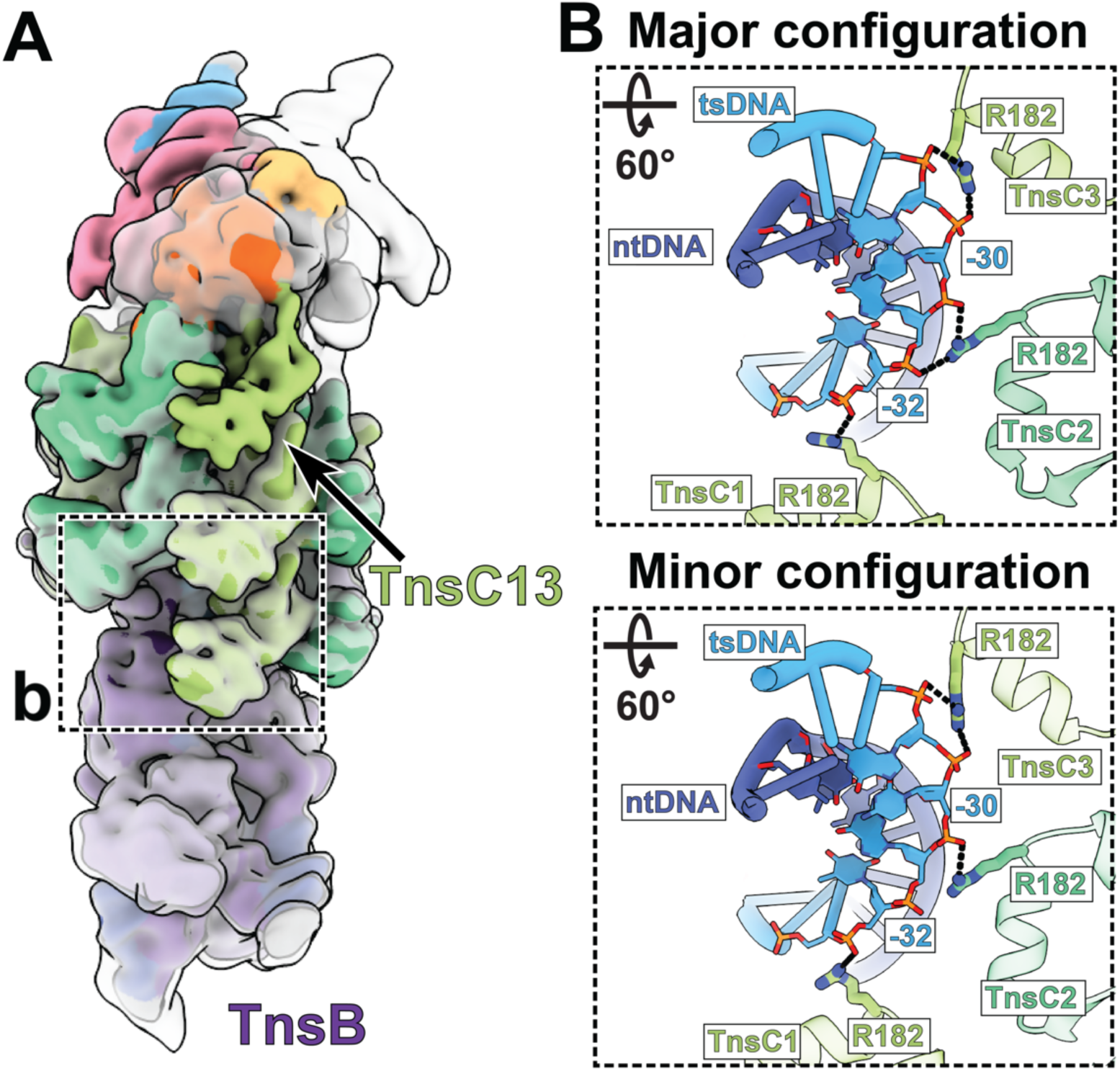
TnsC-DNA interaction proximal to TnsB is identical in the major and minor configurations. **A.** pass filtered (10 Å) cryo-EM reconstructions of both major (transparent grey) and minor (solid, colored) configurations are aligned with respect to TnsB. The dashed box indicates the TnsC-DNA interactions at the TnsB proximal region shown as inset in panel B. **B**. Three TnsB-proximal TnsC protomers (From TnsC1 to TnsC3) in both major and minor configurations interact with target strand DNA (tsDNA) in an identical manner through residue R182. Hydrogen bonding interactions (distance cut off <4Å) between protein residues and the sugar-phosphate backbone of DNA are represented with dashed lines. Base pairs that are interacting with TnsC are represented as filled nucleotides and stick phosphate-backbone.

**Table S1:** Cryo-EM Map and Model Validation

| Name | TnsB<sup>CTD</sup>-TnsC-TniQ complex | Transpososome major configuration | Transpososome minor configuration |
| --- | --- | --- | --- |
| PDB ID | 7SVU | 8EA3 | 8EA4 |
| EMDB ID | EMD-25453 | EMD-27971 | EMD- 27972 |
| **Data collection and Processing** |  |  |  |
| Microscope | Talos-Arctica | Titan-Krios | Titan-Krios |
| Voltage (kV) | 200 | 300 | 300 |
| Camera | K3 | K3 | K3 |
| Magnification | 63,000 | 81,000 | 81,000 |
| Pixel size at detector (Å/pixel) | 1.330 | 1.067 | 1.067 |
| Total electron exposure (e-/Å<sup>2</sup>) | 50.00 | 49.91 | 49.91 |
| Exposure rate (e-/pixel/sec) | 28.0 | 21.9 | 21.9 |
| Number of frames | 50 | 50 | 50 |
| Defocus range (µm) | -1 – -2.5 | -0.8 – -2.5 | -0.8 – -2.5 |
| Automation software | SerialEM | Leginon | Leginon |
| Energy filter slit width | 20 eV | 20 eV | 20 eV |
| Micrographs collected (no.) | 1,271 | 17,554 | 17,554 |
| Micrographs used (no.) | 1,263 | 14,017 | 14,017 |
| Total extracted particles (no.) | 795,631 | 535,932 | 535,932 |
| **For each reconstruction:** |  |  |  |
| Refined particles (no.) | 624,597 | 285,090 | 250,842 |
| Final particles (no.) | 61,515 | 188,055 | 67,096 |
| Symmetry | C1 | C1 | C1 |
| Resolution (global, Å) |  |  |  |
| FSC 0.5 (unmasked/masked) | 7.4/4.0 | 8.8/3.9 | 9.6/4.2 |
| FSC 0.143 (unmasked/masked) | 4.1/3.5 | 6.9/3.5 | 7.2/3.7 |
| Resolution range (local, Å) | 3.0 – 7.0 | 3.0 – 7.0 | 3.0 – 7.0 |
| Resolution range due to anisotropy (Å) | 3.2 – 3.7 | 3.6 – 4.1 | 3.8 – 4.4 |
| Map sharpening *B* factor (Å<sup>2</sup>) | - | 85.3 | 70.7 |
| Map sharpening methods | LocSpiral | cryoSPARC | cryoSPARC |
| **Model composition** |  |  |  |
| Protein residues | 3,283 | 5,618 | 5,875 |
| Ligands | 22 | 26 | 28 |
| RNA/DNA | 52 | 495 | 495 |
| **Model Refinement** |  |  |  |
| Refinement package | Coot/Rosetta/Phenix | Coot/Rosetta/Phenix | Coot/Rosetta/Phenix |
| - real or reciprocal space | Real space | Real space | Real space |
| - resolution cutoff (Å) | 3.5 | 3.2 | 3.8 |
| Model-Map scores |  |  |  |
| - CC (box) | 0.80 | 0.79 | 0.77 |
| - Average FSC (0.5 cutoff, Å) | 3.89 | 4.11 | 4.42 |
| R.m.s deviations from ideal values |  |  |  |
| Bond length (Å) | 1.826 | 1.714 | 1.665 |
| Bond angles (°) | 0.019 | 0.012 | 0.012 |
| **Validation** |  |  |  |
| MolProbity score | 1.75 | 1.68 | 1.79 |
| CaBLAM outliers (%) | 1.38 | 1.15 | 1.10 |
| Clashscore | 14.46 | 10.95 | 15.13 |
| Poor rotamers (%) | 0.42 | 0.33 | 0.23 |
| C-beta outliers (%) | 0.39 | 0.25 | 0.22 |
| EMRinger score | 1.75 | 1.50 | 1.19 |
| Ramachandran plot |  |  |  |
| Favored (%) | 97.59 | 97.34 | 97.47 |
| Outliers (%) | 0.15 | 0.18 | 0.15 |

**Table S2.**
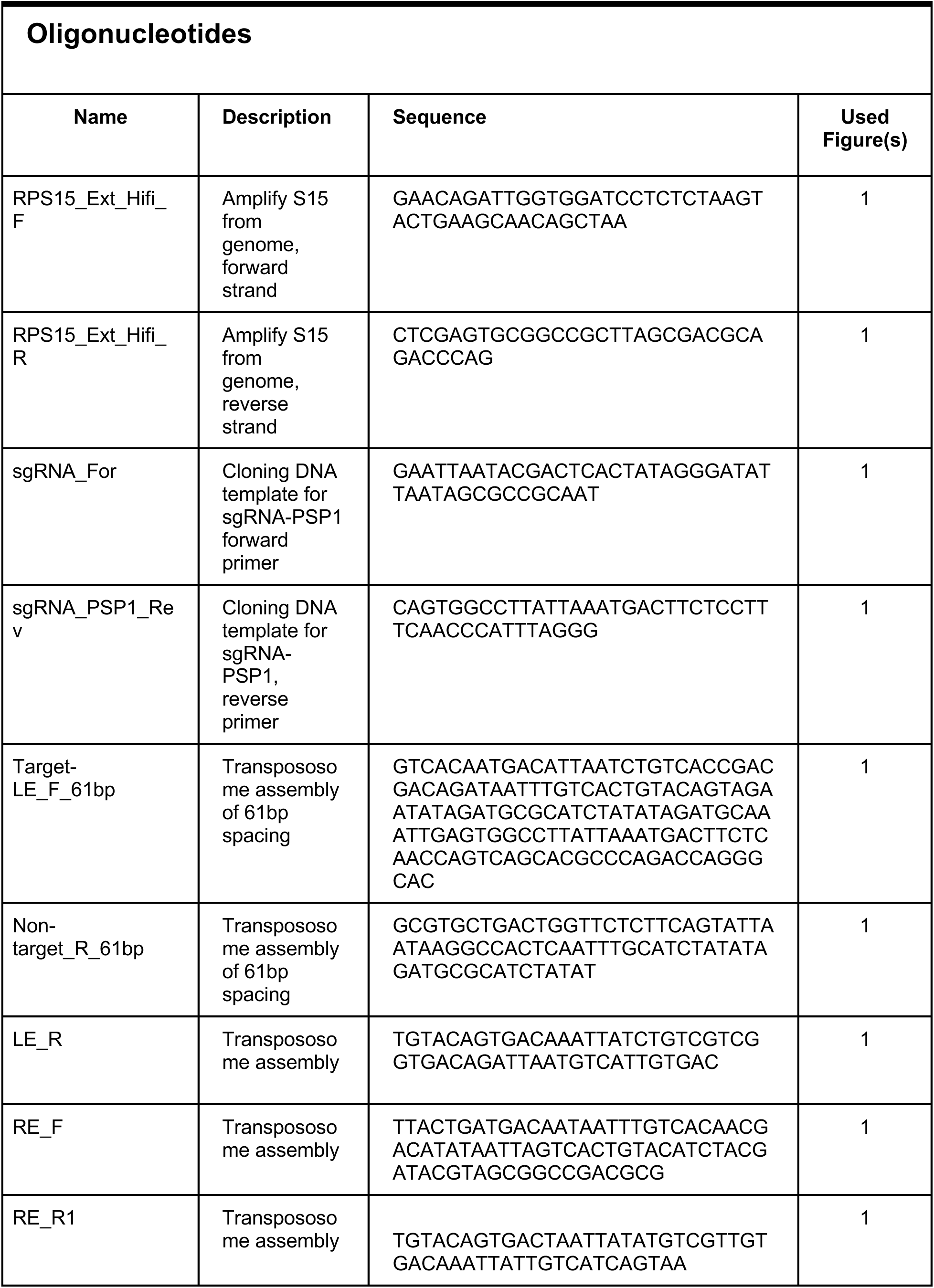
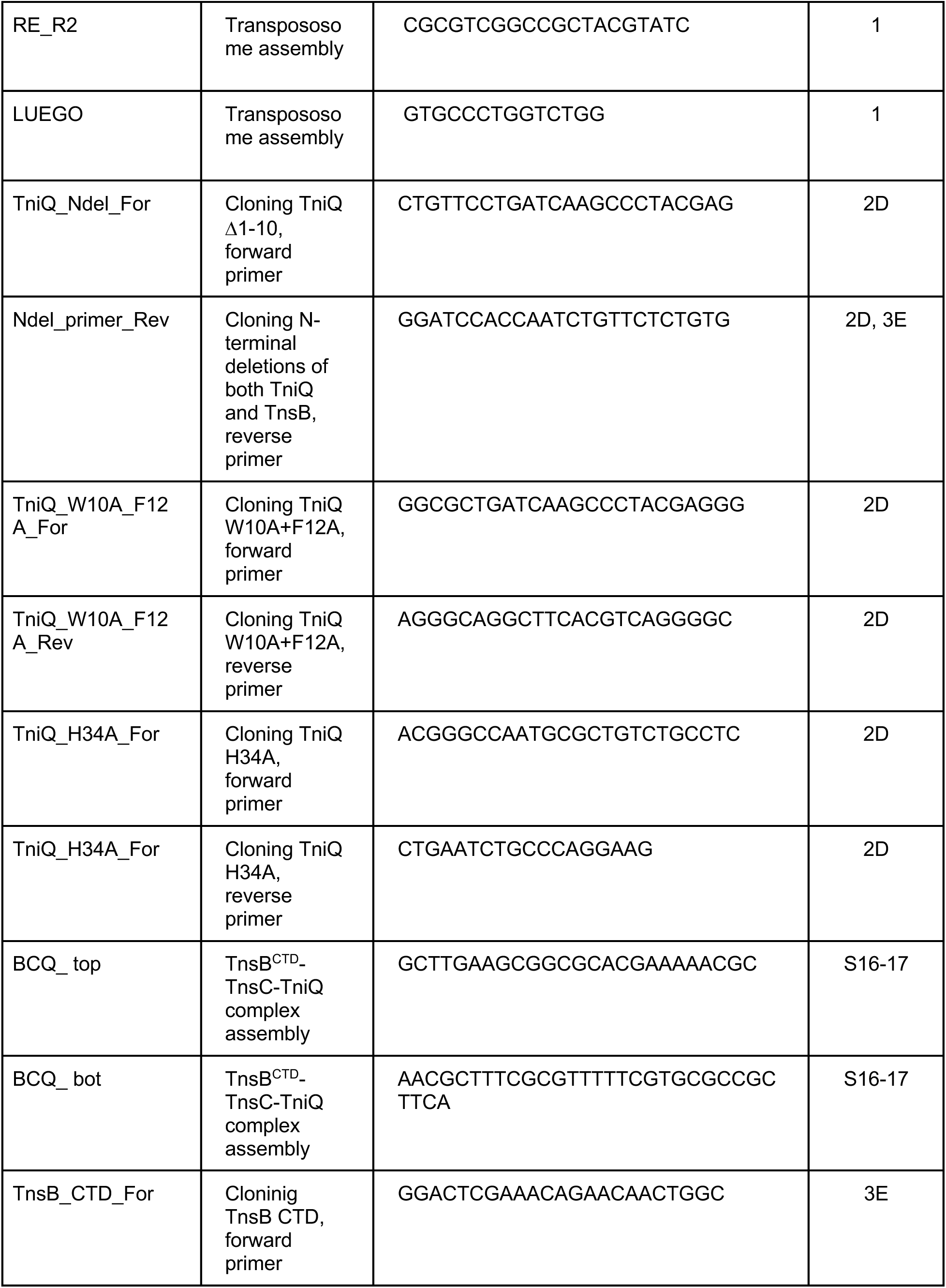
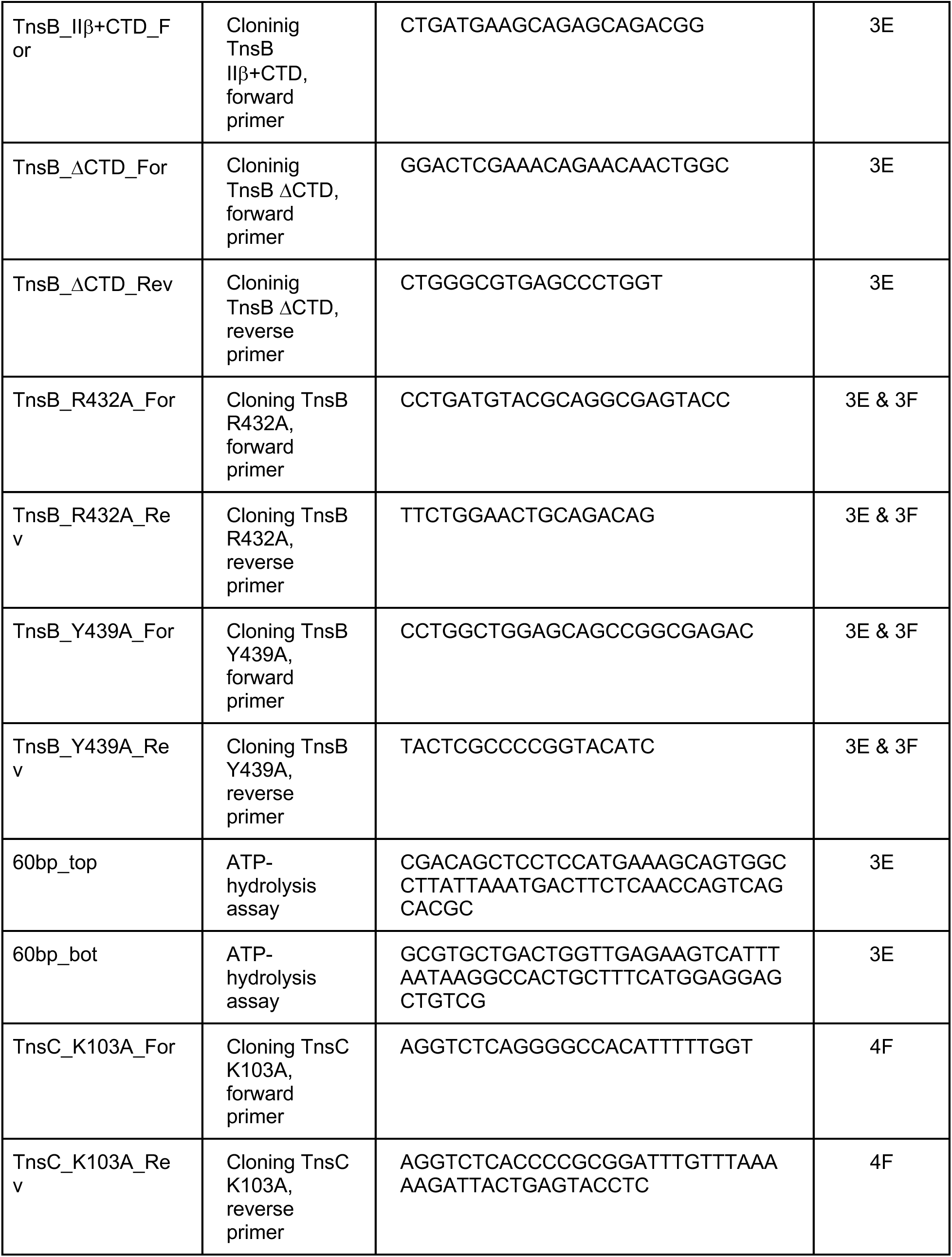
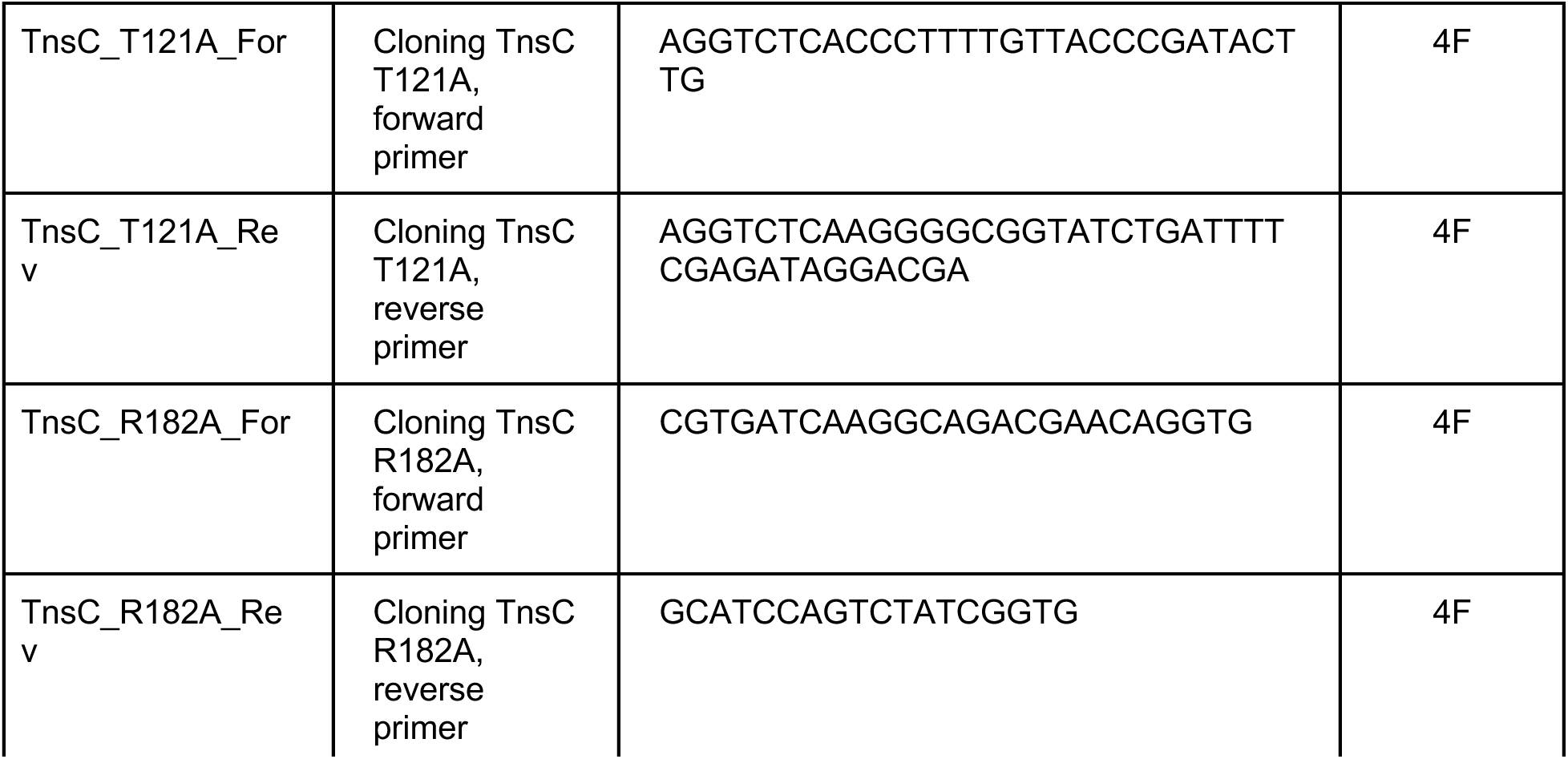
Oligonucleotides used in this study

**Table S3.** The distance between TnsC residues and the closest atom from the phosphate backbone of DNA. Distances were measured between potential hydrogen-bonding donor atoms and the closest oxygen atom from the DNA phosphate backbone. Most of the Lys103 residues (11 out of 12) are too far from the DNA backbone to form a hydrogen bonding interaction. Thr121 and Arg182 interact with the non-target strand (nt) or the target-strand (ts) throughout the transpososome. Lys150 contacts the non-target strand near the Cas12k of the transpososome. Lys119 residues are positioned following the target strand, but most of the residues (7 out of 12, indicated with red asterisks) do not have a strong cryo-EM density of the side chain. These residues were built with the most common rotamers for measuring the distance. Black asterisks indicate the uncertainty of the measured distances due to the lack of side-chain density. The closest strand is annotated only when the distance is less than or equal to 4Å, and annotated as a dot (.) otherwise.

|  | Lys103 |  | Thr121 |  | Lys150 |  | Arg182 |  | Lys119* |  |
| --- | --- | --- | --- | --- | --- | --- | --- | --- | --- | --- |
| TnsC | DNA | Distance | DNA | Distance | DNA | Distance | DNA | Distance | DNA | Distance |
| 12 | . | 4.8 Å | nt | 2.2 Å | nt | 3.1 Å | . | 5.7 Å | . | 4.6 Å |
| 11 | ts | 3.9 Å | ts | 3.6 Å | nt | 3.9 Å | . | 5.4 Å | . | 5.1 Å |
| 10 | . | 4.2 Å | . | 4.5 Å | . | 4.5 Å | . | 4.6 Å | . | 6.4 Å* |
| 9 | . | 4.4 Å | . | 5.1 Å | . | 4.7 Å | . | 6.0 Å | . | 4.5 Å* |
| 8 | . | 4.6 Å | . | 5.0 Å | . | 5.1 Å | . | 6.2 Å | . | 4.6 Å* |
| 7 | . | 5.3 Å | . | 5.2 Å | . | 4.9 Å | . | 5.5 Å | . | 4.4 Å* |
| 6 | . | 5.4 Å | . | 5.7 Å | . | 4.6 Å | . | 5.6 Å | ts | 3.1 Å* |
| 5 | . | 4.6 Å | . | 4.3 Å | . | 5.7 Å | . | 4.6 Å | . | 4.3 Å |
| 4 | . | 4.9 Å | nt | 4.0 Å | . | 7.1 Å | . | 5.0 Å | ts | 3.9 Å |
| 3 | . | 4.1 Å | nt | 2.5 Å | . | 5.6 Å | ts | 2.7 Å | ts | 3.1 Å* |
| 2 | . | 4.3 Å | nt | 2.2 Å | . | 4.7 Å | ts | 3.6 Å | ts | 2.6 Å* |
| 1 | . | 5.9 Å | nt | 2.5 Å | . | 4.9 Å | ts | 3.3 Å | . | 4.3 Å |

## References

1. J. E. Peters, K. S. Makarova, S. Shmakov, E. V. Koonin, Recruitment of CRISPR-Cas systems by Tn7-like transposons. Proc Natl Acad Sci U S A 114, E7358–E7366 (2017).

2. J. Strecker et al., RNA-guided DNA insertion with CRISPR-associated transposases. Science 365, 48–53 (2019).

3. S. E. Klompe, P. L. H. Vo, T. S. Halpin-Healy, S. H. Sternberg, Transposon-encoded CRISPR-Cas systems direct RNA-guided DNA integration. Nature 571, 219–225 (2019).

4. G. Faure et al., CRISPR-Cas in mobile genetic elements: counter-defence and beyond. Nat Rev Microbiol 17, 513–525 (2019).

5. M. T. Petassi, S. C. Hsieh, J. E. Peters, Guide RNA Categorization Enables Target Site Choice in Tn7-CRISPR-Cas Transposons. Cell 183, 1757–1771 e1718 (2020).

6. M. Saito et al., Dual modes of CRISPR-associated transposon homing. Cell, (2021).

7. S. Benler et al., Cargo Genes of Tn7-Like Transposons Comprise an Enormous Diversity of Defense Systems, Mobile Genetic Elements, and Antibiotic Resistance Genes. mBio 12, e0293821 (2021).

8. J. E. Peters, Targeted transposition with Tn7 elements: safe sites, mobile plasmids, CRISPR/Cas and beyond. Mol Microbiol 112, 1635–1644 (2019).

9. T. S. Halpin-Healy, S. E. Klompe, S. H. Sternberg, I. S. Fernández, Structural basis of DNA targeting by a transposon-encoded CRISPR-Cas system. Nature 577, 271–274 (2020).

10. M. Schmitz, I. Querques, S. Oberli, C. Chanez, M. Jinek, Structural basis for RNA-mediated assembly of type V CRISPR-associated transposons. bioRxiv, (2022).

11. P. L. Sharpe, N. L. Craig, Host proteins can stimulate Tn7 transposition: a novel role for the ribosomal protein L29 and the acyl carrier protein. EMBO J 17, 5822–5831 (1998).

12. F. T. Hoffmann et al., Selective TnsC recruitment enhances the fidelity of RNA-guided transposition. Nature, (2022).

13. K. Y. Choi, J. M. Spencer, N. L. Craig, The Tn7 transposition regulator TnsC interacts with the transposase subunit TnsB and target selector TnsD. Proc. Natl. Acad. Sci. U. S. A. 111, E2858–2865 (2014).

14. C. S. Waddell, N. L. Craig, Tn7 transposition: two transposition pathways directed by five Tn7-encoded genes. Genes Dev 2, 137–149 (1988).

15. S. E. Klompe et al., Evolutionary and mechanistic diversity of Type I-F CRISPR-associated transposons. Mol Cell 82, 616–628 e615 (2022).

16. I. Querques, M. Schmitz, S. Oberli, C. Chanez, M. Jinek, Target site selection and remodelling by type V CRISPR-transposon systems. Nature, (2021).

17. R. Xiao et al., Structural basis of target DNA recognition by CRISPR-Cas12k for RNA-guided DNA transposition. Mol Cell, (2021).

18. J. U. Park, A. W. Tsai, T. H. Chen, J. E. Peters, E. H. Kellogg, Mechanistic details of CRISPR-associated transposon recruitment and integration revealed by cryo-EM. Proc Natl Acad Sci U S A 119, e2202590119 (2022).

19. J. U. Park et al., Structural basis for target site selection in RNA-guided DNA transposition systems. Science 373, 768–774 (2021).

20. Y. Shen et al., Structural basis for DNA targeting by the Tn7 transposon. Nat Struct Mol Biol 29, 143–151 (2022).

21. P. A. Gary, M. C. Biery, R. J. Bainton, N. L. Craig, Multiple DNA processing reactions underlie Tn7 transposition. J Mol Biol 257, 301–316 (1996).

22. S. P. Montano, Y. Z. Pigli, P. A. Rice, The mu transpososome structure sheds light on DDE recombinase evolution. Nature 491, 413–417 (2012).

23. N. Jullien, J. P. Herman, LUEGO: a cost and time saving gel shift procedure. Biotechniques 51, 267–269 (2011).

24. F. Wang et al., General and robust covalently linked graphene oxide affinity grids for high-resolution cryo-EM. Proc Natl Acad Sci U S A 117, 24269–24273 (2020).

25. A. Patel, D. Toso, A. Litvak, E. Nogales, Efficient graphene oxide coating improves cryo-EM sample preparation and data collection from tilted grids. bioRxiv, (2021).

26. C. Suloway et al., Automated molecular microscopy: the new Leginon system. J Struct Biol 151, 41–60 (2005).

27. S. Q. Zheng et al., MotionCor2: anisotropic correction of beam-induced motion for improved cryo-electron microscopy. Nat Methods 14, 331–332 (2017).

28. G. C. Lander et al., Appion: an integrated, database-driven pipeline to facilitate EM image processing. J Struct Biol 166, 95–102 (2009).

29. A. Punjani, J. L. Rubinstein, D. J. Fleet, M. A. Brubaker, cryoSPARC: algorithms for rapid unsupervised cryo-EM structure determination. Nat Methods 14, 290–296 (2017).

30. S. H. Scheres, RELION: implementation of a Bayesian approach to cryo-EM structure determination. J Struct Biol 180, 519–530 (2012).

31. S. H. Scheres, Processing of Structurally Heterogeneous Cryo-EM Data in RELION. Methods Enzymol 579, 125–157 (2016).

32. J. Zivanov et al., New tools for automated high-resolution cryo-EM structure determination in RELION-3. Elife 7, (2018).

33. J. Zivanov, T. Nakane, S. H. W. Scheres, A Bayesian approach to beam-induced motion correction in cryo-EM single-particle analysis. IUCrJ 6, 5–17 (2019).

34. E. F. Pettersen et al., UCSF Chimera--a visualization system for exploratory research and analysis. J Comput Chem 25, 1605–1612 (2004).

35. A. Punjani, D. J. Fleet, 3D variability analysis: Resolving continuous flexibility and discrete heterogeneity from single particle cryo-EM. J Struct Biol 213, 107702 (2021).

36. P. Emsley, B. Lohkamp, W. G. Scott, K. Cowtan, Features and development of Coot. Acta Crystallogr D Biol Crystallogr 66, 486–501 (2010).

37. J. Jumper et al., Highly accurate protein structure prediction with AlphaFold. Nature 596, 583–589 (2021).

38. P. V. Afonine et al., New tools for the analysis and validation of cryo-EM maps and atomic models. Acta Crystallogr D Struct Biol 74, 814–840 (2018).

39. T. D. Goddard et al., UCSF ChimeraX: Meeting modern challenges in visualization and analysis. Protein Sci 27, 14–25 (2018).

40. C. J. Williams et al., MolProbity: More and better reference data for improved all-atom structure validation. Protein Sci 27, 293–315 (2018).

41. D. N. Mastronarde, Automated electron microscope tomography using robust prediction of specimen movements. J Struct Biol 152, 36–51 (2005).

42. D. Tegunov, P. Cramer, Real-time cryo-electron microscopy data preprocessing with Warp. Nat Methods 16, 1146–1152 (2019).

43. T. Bepler et al., Positive-unlabeled convolutional neural networks for particle picking in cryo-electron micrographs. Nat Methods 16, 1153–1160 (2019).

44. S. Kaur et al., Local computational methods to improve the interpretability and analysis of cryo-EM maps. Nat Commun 12, 1240 (2021).

45. W. M. Cianfrocco MA, Youn C, Wagner R, Leschziner AE., COSMIC²: A Science Gateway for Cryo-Electron Microscopy Structure Determination. Practice & Experience in Advanced Research Computing, (2017).

46. A. Leaver-Fay et al., ROSETTA3: an object-oriented software suite for the simulation and design of macromolecules. Methods Enzymol 487, 545–574 (2011).

